# Toughening mechanisms for the attachment of architectured materials: The mechanics of the tendon enthesis

**DOI:** 10.1101/2021.05.17.444505

**Authors:** Mikhail Golman, Adam C. Abraham, Iden Kurtaliaj, Brittany P. Marshall, Yizhong Jenny Hu, Andrea G. Schwartz, X. Edward Guo, Victor Birman, Philipp J. Thurner, Guy M. Genin, Stavros Thomopoulos

## Abstract

Architectured materials offer tailored mechanical properties but are limited in engineering applications due to challenges in maintaining toughness across their attachments. The enthesis connects tendon and bone, two vastly different architectured materials, and exhibits toughness across a wide range of loadings. Understanding the mechanisms by which this is achieved could inform the development of engineered attachments. Integrating experiments, simulations, and novel imaging that enabled simultaneous observation of mineralized and unmineralized tissues, we identified putative mechanisms of enthesis toughening in a mouse model and then manipulated these mechanisms via *in vivo* control of mineralization and architecture. Imaging uncovered a fibrous architecture within the enthesis that controls trade-offs between strength and toughness. *In vivo* models of pathology revealed architectural adaptations that optimize these trade-offs through cross-scale mechanisms including nanoscale protein denaturation, milliscale load-sharing, and macroscale energy absorption. Results suggest strategies for optimizing architecture for tough bimaterial attachments in medicine and engineering.

**Teaser:** The architecture of the tendon-to-bone attachment is designed for toughness.

## Introduction

Materials whose micro- and meso-scale architectures endow them with useful mechanical functions are found throughout nature and, more recently, in engineering (*1–4*). However, engineering application of such architectured materials is limited by the challenge of attaching them (*5*). Typical features of architectured materials (e.g., microtruss composites) lead to local elevations in stress that can reduce strength (i.e., stress required to break the material) and toughness (i.e., the energy absorbed prior to breaking the material) when they are connected to other materials (*6, 7*). Natural materials provide a rich source of inspiration for the design and attachment of architectured engineering materials. For example, the tendon enthesis illustrates a number of novel and often counter-intuitive ways by which architectured materials can be effectively connected. Tendon and bone, tissues with a two orders-of-magnitude difference in modulus, display a hierarchical architecture ranging from nanometer-scale triple-helix tropocollagen molecules to sub-micrometer-diameter fibrils to 10-100 micrometer-diameter fibers that extend over millimeters (*8*). Across species, strong attachment of tendon and bone arises from a zone of compliant transitional tissue (*9, 10*) that mitigates stress concentrations through allometric scaling of geometry (*11*) and through functional gradations of both fiber orientation (*12, 13*) and bioapatite mineral (*14–17*). These aspects of enthesis architecture are not recreated following injury, and surgical repairs thus often fail (*18, 19*). Despite progress in understanding how the enthesis achieves a strong attachment under sub-damage loading regimes, it remains unclear how toughness is achieved to prevent interfacial failure. Understanding these mechanisms will guide engineering and medical approaches to bimaterial attachment.

We therefore aimed to identify enthesis architectural and compositional toughening mechanisms in mice using imaging, biomechanical testing, and mathematical modeling. A novel micro computed tomography (microCT) technique was developed to simultaneously visualize the mineralized and unmineralized fibrous networks within the tendon-bone attachment at sub-micrometer resolution. We manipulated the fibrous network through pathophysiologic loading *in vivo* in a mouse model and quantified how monotonic (acute) and cyclical (degenerative) loading affected enthesis strength and toughness. Biomechanical analysis and numerical simulation supported our hypothesis that architectural toughening arises from the composition (nanoscale mineral and proteoglycans), structure (microscale collagen organization and recruitment), and position (macroscale loading angle) of the transitional material. Physiologically, enthesis composition and micro-structure *in vivo* adapted to loading in a way that revealed a trade-off between strength and toughness. These features of the adaptable, architectured, fibrous enthesis have direct implications for tough attachment between dissimilar materials, facilitating improved design of surgical and tissue engineered solutions for tendon-to-bone repair.

## Results

### Attachment at the enthesis relies on a fibrous architectured material system

Using microCT imaging with mercury (II) chloride staining, we obtained simultaneous, sub-micrometer imaging of unmineralized and mineralized tissue in the mouse supraspinatus tendon enthesis (Fig. 1A) and discovered that the function of the enthesis had been previously misunderstood. Hidden within the well-known attachment footprint (*11*) (Fig. 1B within blue dotted line) was a smaller, denser “primary” insertion site where tendon fibers directly inserted into bone over 30% ± 3.5% of the footprint area (Fig. 1C, within green dotted line, FigS1, Movie S1). Collagen fibers were continuous from muscle to bone but branched into smaller diameter fibers as they inserted into bone on one end, as previously described (*9, 20*), or muscle at the other.

**Fig. 1.**
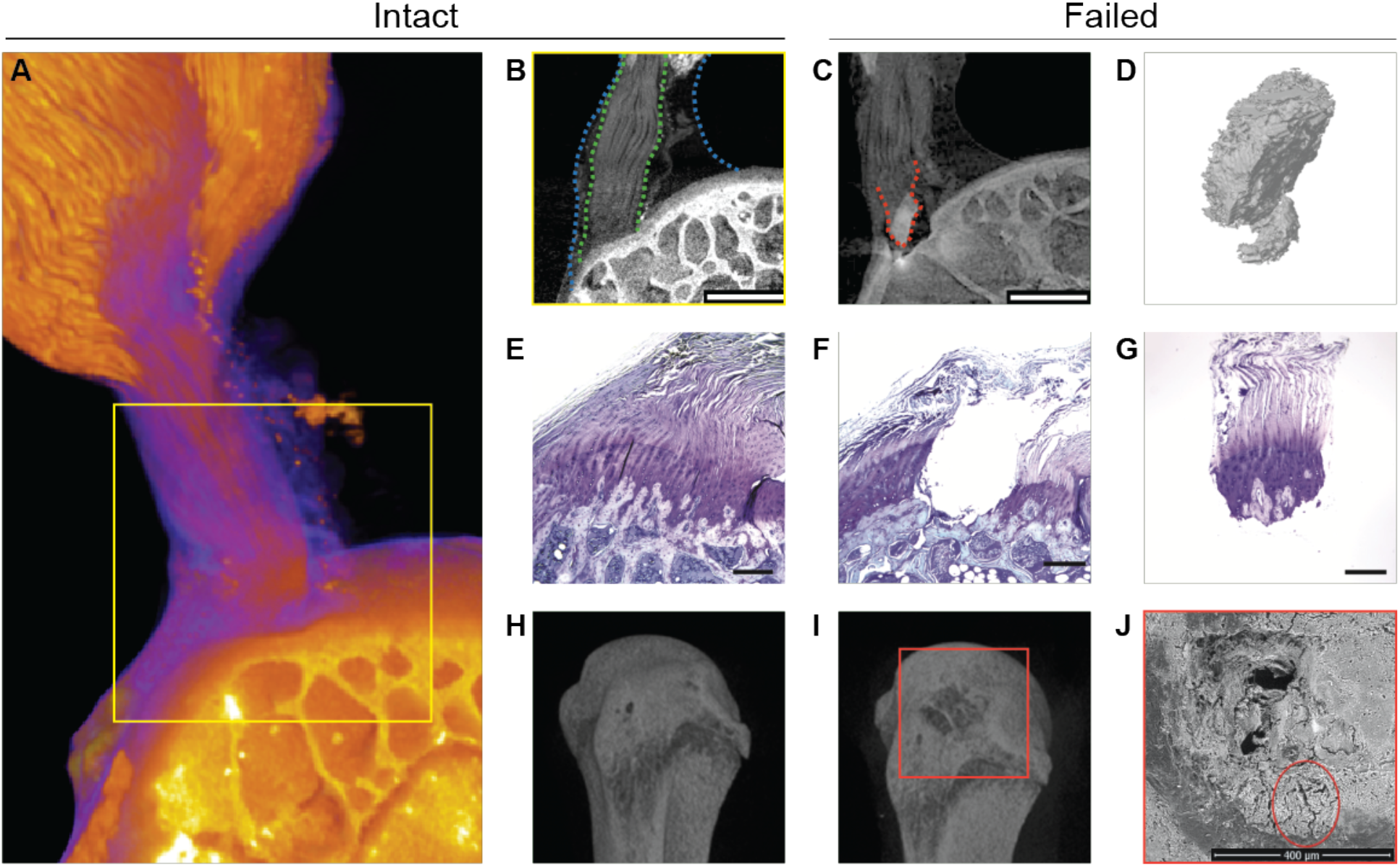
The tendon enthesis exhibits a fibrous architectured material system that fails via bony avulsion under quasi-static loading. **(A)-(C),** Mercury (II) chloride-stained contrast enhanced high-resolution microCT imaging revealed that, hidden within the well-known larger apparent attachment footprint area, is a smaller, much denser primary insertion site where tendon fibers insert directly into the bone. Imaging revealed that, under quasi-static loading, only 47.4+/-5.1% of the apparent attachment site was avulsed, revealing a previously unknown primary attachment. (A) Three-dimensional volume rendering of representative intact enthesis. (B) Magnified cross sectional view of yellow box in a; within blue dotted lines outline apparent enthesis and within green dotted lines outline dense primary insertion. (scale: 500 μm). (C) Post-failure imaging showing avulsed bony fragment at primary insertion site, outlined with a red dotted line. (scale: 500 μm). **(D)** Three-dimensional representation of avulsed fragment showing portions of trabeculae at the failure site. **(E)-(G)**, Histological sections of (E) intact, and (F)-(G) failed enthesis stained with toluidine blue (scale: 250 μm). **(H)-(I)**, Three-dimensional reconstruction from conventional microCT imaging of a representative (H) intact and (I) failed enthesis sample. **(J)** Scanning electron microscopy of the failure site showing crack propagation around the avulsion site, outlined by a red circle (scale 400 μm).

To test the hypothesis that the primary insertion site was responsible for load transfer, we stretched supraspinatus tendon enthesis specimens to failure quasi-statically. The enthesis failed through avulsion of a bone plug (Fig. 1C) over 22.4% ± 6.2% (0.31 ± 0.09 mm^2^) of the apparent insertion site (Fig.S1C), with the majority of the primary insertion avulsed, but with peritenon tissue surrounding the primary insertion site still attached (Fig. S2A, Movie S2). Failure occurred catastrophically, with little resistance to post-failure force (Fig. S2B), supporting our hypothesis.

We next asked how the primary insertion resisted failure loads. Although failure was expected at the mineralized interface within fibrocartilage where the stress concentrations were predicted to occur (*21, 22*), this was not observed, indicating mechanisms to alleviate these stress concentrations. Failure occurred either at the interface between mineralized fibrocartilage and bone (MF-B failure type), or within trabecular bone (B-T failure type) (Fig. 1 C - I; Fig. S3A, Movie S3) and in all cases with crack propagation around the avulsion site (scanning electron microscopy, Fig. 1J and Fig. 3SB). For this loading, the fibrous primary enthesis was thus tougher than cortical bone, with the more compliant fibrocartilage storing enough energy to fracture and avulse bone.

### Multiscale toughening mechanisms enable resistance to cyclical loading

The enthesis is durable against the complex and repeated loadings of daily activities (*23*), but failure mechanisms change with loading regime and age. Avulsions are common in high-impact injuries for pediatric patients (*24*), but rupture at the tendon end of enthesis is prominent in degenerated rotator cuffs of adult patients (*25–27*). We therefore hypothesized that toughening mechanisms depend upon the loading regime.

In response to acute loading (monotonic tension across a range of loading rates) or fatigue loading (cyclic loading at 2 Hz, either 1-20 % or 20-70% failure load), three distinct failure modes were observed (Fig. 2 A-E): bone avulsion, tendon mid-substance failure, and tendon-bone interface failure. Acute loading led primarily to avulsion, regardless of loading rate. Although enthesis mechanical properties were largely strain-rate insensitive, like tendon properties (*28, 29*), strength (failure load) and toughness (work to failure, calculated as the area under the force-displacement curve) increased at higher strain rates by as much as 1.4-fold (*p*<0.0001) and 1.6-fold (*p*<0.01), respectively, compared to that of control test case (*n*=10-12/case, Fig. 2 C and D; Fig.S4 A-C). Notably, the area and number of fragments of the avulsed region increased with loading rate (Fig. S4 C-F). In contrast to acute loading, all cyclically loaded samples (High: 2 Hz, 20-70% of strength) failed in the unmineralized fibrocartilage portion of the attachment (“insertion failure”, Fig 2E). Samples cyclically loaded at lower, physiologically relevant loads (Low: 2Hz, 1%-20% of strength), did not fail, even after 100,000 cycles (Fig. 2E), indicating that these loading levels were under the enthesis fatigue limit. Results thus suggested that the mechanisms protecting fibrocartilaginous enthesis tissue might be gradually attenuated under sufficiently severe cyclical loading.

**Fig. 2.**
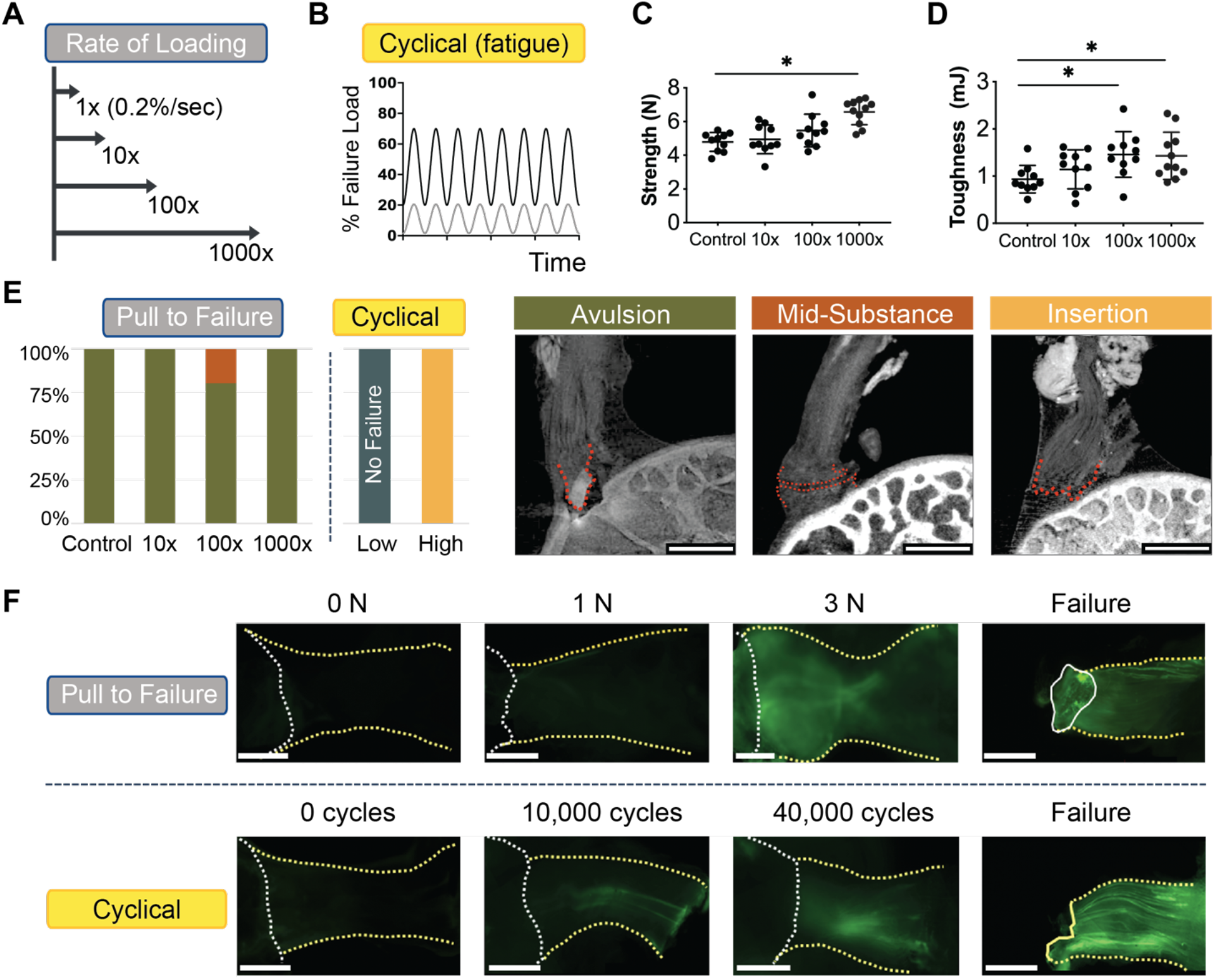
Multiscale toughening mechanisms enable the entheses to exhibit distinct failure modes under varying loading conditions. **(A)-(B)**, To examine effect of loading on failure mode, samples were loaded (A) across a range of loading rates to simulate acute injuries or (B) loaded cyclically to simulate degenerative loading. **(C)** Enthesis strength (i.e., failure load) and **(D)** enthesis toughness (i.e., energy absorption) increased with the loading rate. (**** p<0.0001, * p<0.05, ANOVA followed by the Dunnet’s multiple comparison test). **(E)** There were three distinct failure modes, depending on the loading regime: bone avulsion, tendon mid-substance, and tendon-bone interface (insertion failure) (scale: 500 μm). Under monotonic loading, most samples failed by bony avulsion failures. Under “high” cyclical loading (20%-70% failure force), all samples failed at the insertion. Under “low” cyclical loading (1%-20% failure force) samples did not fail, even after 100,000 cycles. **(F)** F-CHP fluorescence intensity, indicative of collagen damage accumulation, increased with the level of applied load and with the number of cycles. For quasi-statically loaded samples (F, top), there was little to no fluorescent signal in the low force group (1N-2N), followed by increased staining near the attachment site at higher loads (3N and failure). For cyclically loaded samples (F, bottom), F-CHP staining was initially concentrated in a few fibers near the tendon mid-substance (10K-40K cycles) and ultimately propagated down the entire tendon in concentrated bands (scale: 500 μm).

To identify potential nanoscale mechanisms that could explain this behavior, we quantified molecular damage under the various loading regimes using fluorescein-labeled collagen hybridizing peptide (F-CHP) (*30, 31*). Whole-sample imaging of F-CHP fluorescence intensity, indicative of collagen damage, increased with applied load or number of cycles (Fig. 2F, top). In monotonic loading, fluorescent signal accrued near the primary insertion site when loads exceeded 3 N. Under cyclic loading, signal was concentrated in a few fibers near the tendon mid-substance between 10,000 and 40,000 cycles, then propagated down the entire tendon in concentrated bands (Fig. 2F, bottom). This revealed that, in monotonic loading to failure, energy sufficient to avulse bone was stored in the enthesis with relatively little energy dissipation, while in cyclical loading, energy was absorbed by damage within the tendon and enthesis, eventually leading to failure within the unmineralized tissues. Thus, the enthesis contains fiber-level toughening mechanisms to resist monotonic loading and an underlying nanoscale mechanism to resist cyclical loading.

### Differential recruitment of collagen fibers enables toughness across loading directions

Based upon observations of the fibrous character to the enthesis, we hypothesized that these nanoscale mechanisms are supplemented by macroscale toughening mechanisms to resist failure across a range of directions (i.e., shoulder abduction angle, Fig. 3A-B). Enthesis behavior, including strength and stiffness, varied with the angle of abduction (Fig. 3C-F). This was a surprise given the shoulder’s ability to resist injury across its broad range of motion (*32*). HgCl2-enhanced microCT images revealed that fibers engaged or buckled depending upon loading (Fig. 3B top, Fig.S5A), consistent with fiber recruitment models of tendon mechanics (*33*) and rotator cuff injury (*26*). We therefore developed a series of experiments and models to determine how abduction-dependent fiber architecture and recruitment dictated enthesis mechanics.

**Fig. 3.**
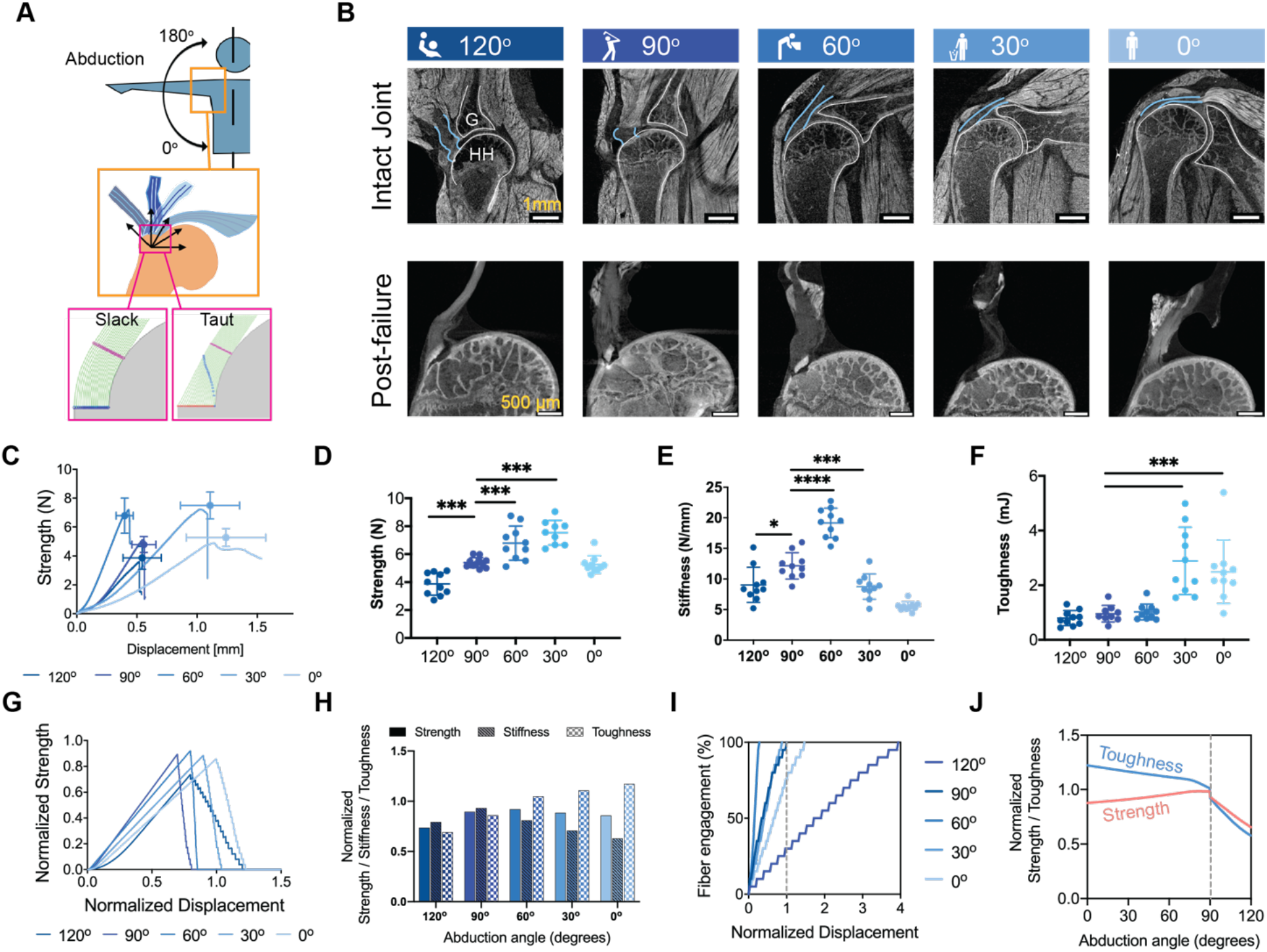
Multiscale toughening mechanisms enable the entheses to exhibit distinct failure modes under varying loading conditions. **(A)** Samples were tested at varying angles of abduction (A, top) and a fiber recruitment model was developed to examine structural and positional contributions to enthesis toughness (A, bottom). **(B)** Contrast-enhanced microCT of intact (b, top row) and failed (b, bottom row) mouse glenohumeral joints at each abduction angle (G: glenoid, HH: humeral head). The supraspinatus tendon (b, top row, outlined in blue) was straight at low abduction angles (0°-30°) and buckled at high abduction angles (90°-120°). **(C)-(F)**, There were significant differences in the attachment mechanical behavior and failure properties when samples were tested quasistatically at varying angles *ex vivo* (C, strength (failure force) vs. displacement plot; D, strength; E, stiffness; F, toughness) (* p<0.05, ** p<0.01, *** p<0.001, **** p<0.0001, ANOVA followed by the Dunnett’s multiple comparison test). **(G)-(J)** A positional recruitment simulation, in which fiber interactions were steric and linear, reproduced experimentally-observed enthesis mechanics as a function of abduction angle. *In silico* (G) strength vs. displacement and (H) strength, stiffness, toughness results normalized against the case when fibers were pulled uniaxially without the geometric constraints. (I) The relationship between fiber engagement and displacement depended on abduction angle, demonstrating that the energy absorbed in re-orienting and engaging fibers drove the toughening behavior of the of attachment. **(J)** Enthesis architecture was optimized for toughness: normalized toughness was generally higher than normalized strength through most abduction angles.

Imaging at 5 μm resolution revealed that the collagen fibers of the supraspinatus tendon enthesis engaged at low abduction angles (0° and 30°) and buckled at high abduction angles (90° and 120°) (Fig. 3B, top row outlined in blue, Video S4). Furthermore, imaging at 0.75 μm resolution confirmed that outer (bursal-side) fibers were longer than inner (articular-side) fibers (Figure 3B, bottom row and Fig.S5A), as similar to what previously described in human (*34, 35*). This indicated that both inner and outer fibers engaged to carry loads at low angles of abduction, but only inner fibers engaged at high angles, with outer fibers remaining slack.

We then explored whether these changes in microscale fiber engagement with shoulder abduction could explain the observed macroscale adaptations in tendon enthesis toughness and strength using a numerical model (Supplemental Text). The model idealized the geometry of the humeral head as a circular bone ridge beneath linear elastic fibers of pre-defined thickness and spacing. Fibers engaged, re-oriented, and contacted neighboring fibers or the humeral head during loading (Fig. 3A bottom, Fig. 5SB) in a way that varied with abduction, and that reproduced trends observed in our experiments (Fig. 3G and H): normalized strength and toughness increased with decreasing abduction angle, while stiffness decreased with decreasing abduction angle. These results thus supported the hypothesis that abduction-dependent fiber recruitment was a factor in failure patterns, with the displacement needed to engage (recruit) all fibers lowest at 60° of abduction (Fig. 3I), and four times higher at 120° than at 90° of abduction. When considering failure behavior across the physiological range of shoulder abduction (Fig. 3J), strength decreased with abduction angle from 90° to 0°, while toughness increased; strength and toughness decreased dramatically beyond 90° of abduction.

From the perspective of shoulder physiology, results inform our understanding of rotator cuff injury. Acute tears in baseball players typically occur in the late-cocking/follow-through phases of pitching (high abduction angles, ~110°) (*36*), consistent with our observations of acute failure via bony avulsion, with size of fractured area lowest at low angles of abduction (*p*< 0.01, FigS6). Rotator cuff tendon tears most commonly initiate on the articular side of the tendon (*25, 26*), consistent with predictions that inner-most fibers engage and fail first at every abduction angle simulated.

The fibrous architecture of the tendon enthesis enabled its fibers to reorient, recruit, and subsequently rupture to balance strength and toughness across a wide range of motion, a tradeoff well known in material design (*7*). The healthy enthesis appeared optimized for toughness, with gains in toughness associated with changing abduction angle achieved through comparably modest losses in strength (Fig 3J). This is somewhat analogous to brittle matrix fibrous composites achieving toughness at the expense of strength (*17, 37*), and how microscale interdigitation of the tendon enthesis toughens attachments (*15*). The trade-off was particularly apparent at lower abduction angles, where rotator cuff muscles were most engaged and enthesis loads were highest (*38*). Although factors such as viscoelasticity and post-yield behavior also contribute to enthesis toughness, the current modeling and experimental results support a clear role for abduction position-dependent kinematics driving tendon enthesis toughness in the rotator cuff.

### Tendon enthesis strength is determined by mineral composition

A spatial gradient in mineral stiffens the enthesis, especially beyond a percolation threshold (*39*), and mitigates stress concentrations (*40*). Proteoglycans stiffen and provide energy dissipation in articular cartilage (*41*). To test the hypothesis that these extracellular matrix components also contribute to enthesis toughness, each was chemically removed from the enthesis prior to mechanical testing (Fig. 4A, Fig.S7). We hypothesized that removal of mineral would reduce stiffness and strength, and that removal of proteoglycans would reduce toughness.

**Fig. 4.**
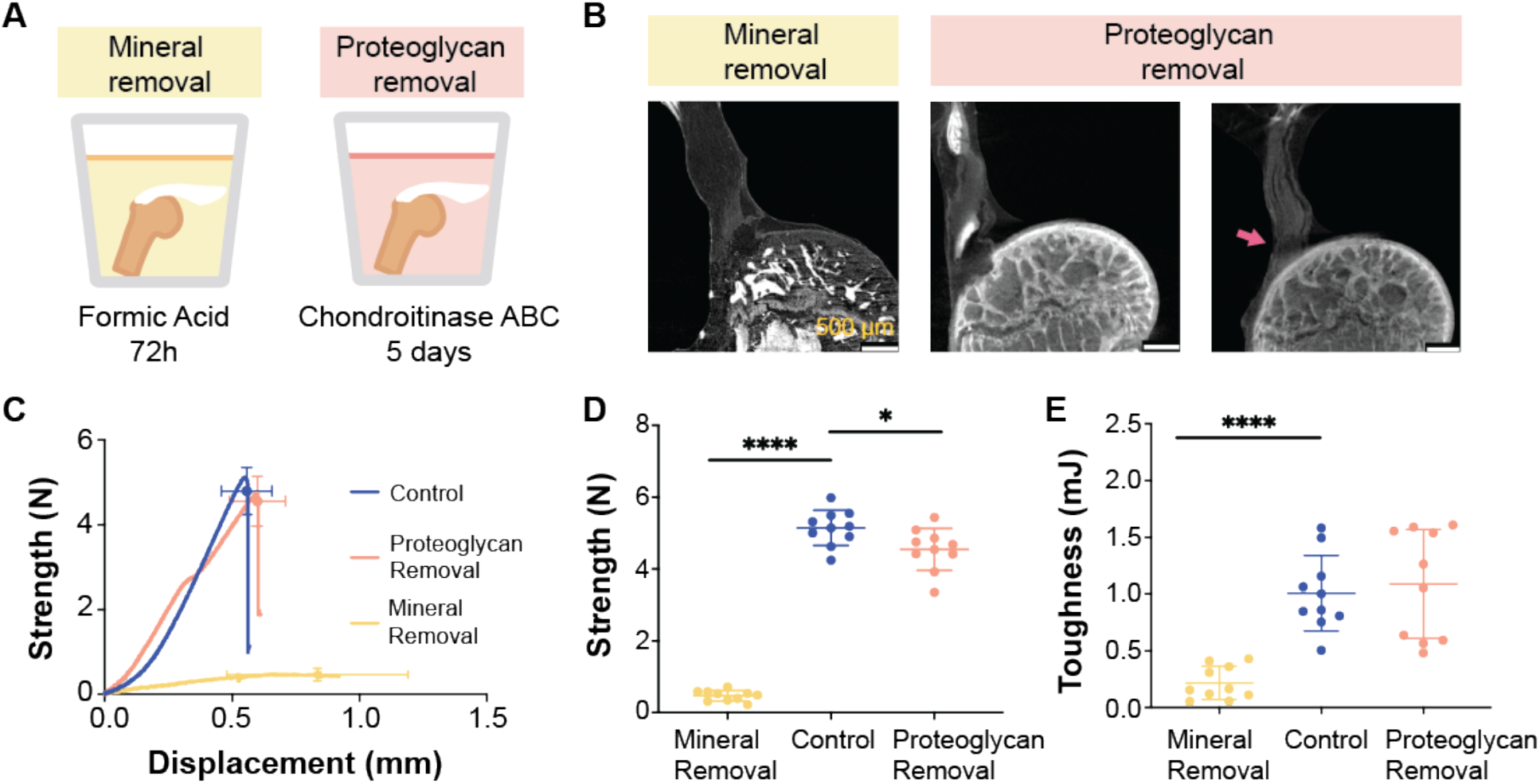
Tendon enthesis composition drives enthesis mechanical properties. **(A)** To examine compositional contributions to tendon-to-bone attachment strength and toughness, samples were immersed in decalcifying agent to completely remove mineral (A, left) or in Chondroitinase ABC for 5 days to chemically digest proteoglycans (A, right). **(B)** Post-failure contrast enhanced microCT scanning showed that loss of mineral or proteoglycan did not significantly alter the failure modes of the tendon enthesis. Most samples failed via bone avulsion, while a small number of samples depleted in proteoglycans failed at the edge of unmineralized fibrocartilage (pink arrow) (scale: 500 μm). **(C)-(E)**, Quasi-static mechanical testing revealed significant differences in mechanical behavior of tendon entheses when mineral was removed. (C) Strength (failure force) vs. displacement behavior. (D) Removal of mineral led to a dramatic decrease in strength; removal of proteoglycan led to a relatively small decrease in strength. (E) Removal of mineral led to a significant decrease in toughness; removal of proteoglycan did not affect enthesis toughness. (* p<0.05, **** p<0.0001, ANOVA followed by the Dunnett’s multiple comparison test).

Removal of mineral or proteoglycan did not significantly alter failure modes under monotonic loading; samples failed primarily via bone avulsion, with 20% (2/10 samples) of mineral depleted samples failing at the insertion (Fig. 4B, Fig. S8A and B). As hypothesized, removal of mineral decreased strength and stiffness (p<0.0001, Fig. 4D and Fig.S8C), but also decreased toughness (p<0.0001, Fig. 4E). Contrary to the hypothesis, removal of proteoglycans did not change toughness, although decreases in strength and stiffness were observed, which were in agreement with prior findings at the scale of collagen fibrils (*42*). Of note, the proteoglycan depletion protocol used here removed proteoglycan in the unmineralized portion of the enthesis only (Fig. S7B), and therefore proteoglycan-mineral interactions cannot be ruled out. Nevertheless, results demonstrate that mineral content is crucial for enthesis strength and toughness.

### The tendon enthesis actively adapts its architecture *in vivo* by controlling mineral composition and microarchitecture

It is well known that bone (*43, 44*) and entheses (*45*) respond to loading by adapting their mineral content. To further elucidate how composition and architecture are modulated at the enthesis *in vivo* to produce toughness, we varied the loading environment of mouse shoulders via botulinum toxin A-induced underuse/paralysis or treadmill-induced overuse (Fig. 5A). We hypothesized that modifications to *in vivo* loading would lead to architectural adaptations that control strength and toughness.

**Fig. 5.**
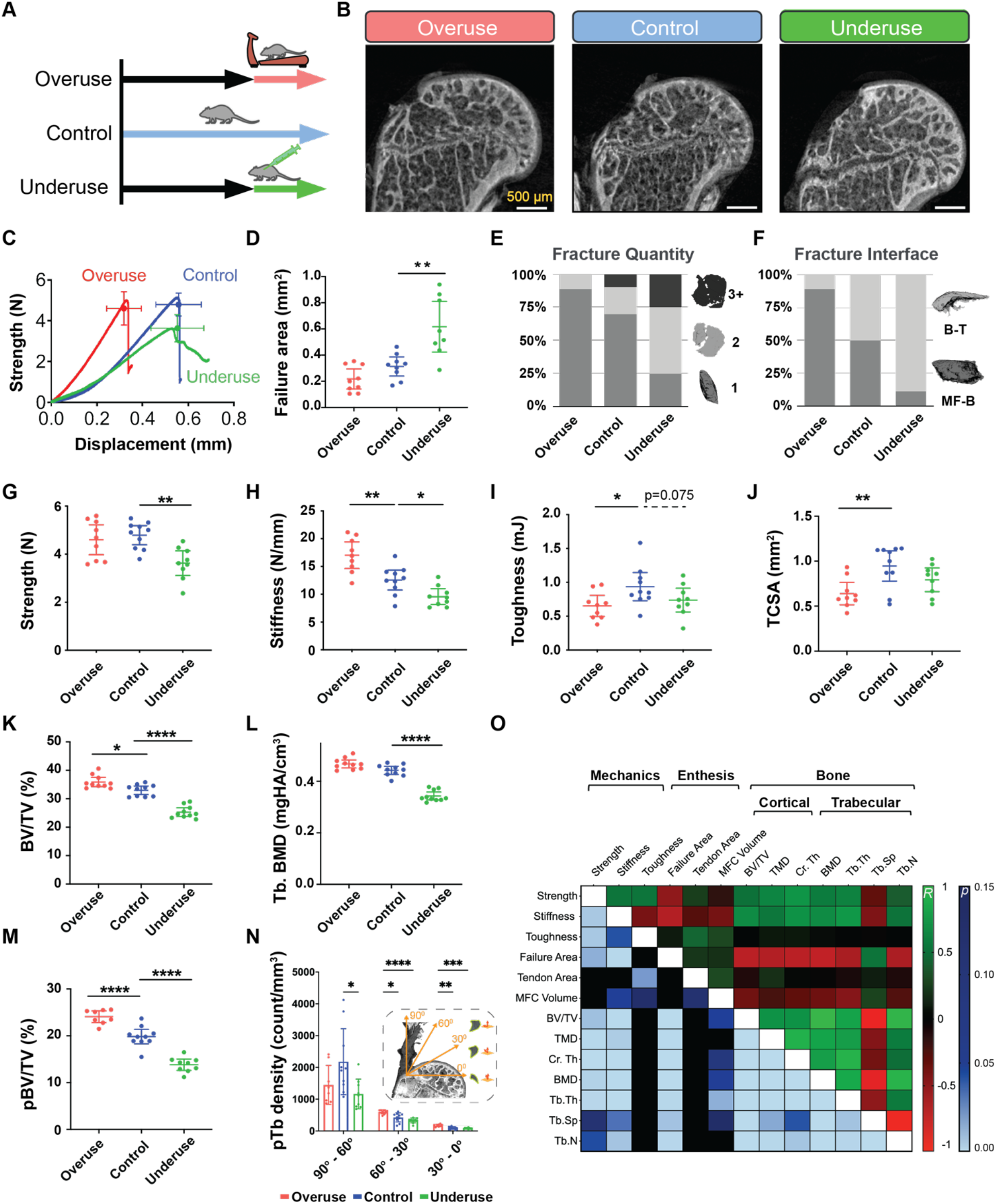
The tendon enthesis actively adapts its architecture in vivo by modifying mineral composition. **(A)** 10-week-old mice were subjected to two degeneration models: underuse degeneration was induced via muscle paralysis and overuse degeneration was achieved through downhill treadmill running for 4 weeks. **(B)** Post-failure contrast enhanced microCT imaging revealed that pathological entheses exhibited exclusively avulsion-type failures under tensile mechanical testing (scale: 500 μm). **(C)-(J)**, Physiological *in vivo* degeneration models reduced the ability of the enthesis to protect against failure. (D) Failure area, (E) avulsed fragment quantity, and (F) failure interfaces were affected by enthesis pathology. Underuse degeneration led to (G) lower strength (p<0.01), (H) lower stiffness (p<0.05), and (I) trended with decreased toughness (p=0.075) compared to that of control. Overuse degeneration decreased (J) tendon cross-sectional area (p<0.01), (H) stiffened the enthesis (p<0.01), and (I) significantly reduced toughness compared to control (p<0.05). **(K)** – **(L)**, Bone morphometric analysis revealed that underuse led to (K) reduced bone volume (BV/TV) (p<0.0001) and (L) reduced bone mineral density (BMD) in the bone underlying the attachment (p<0.0001). **(M)** The volume of load bearing trabecular plates (pBV/TV) increased significantly (p<0.0001) due to overuse and decreased significantly (p<0.0001) due to underuse, with significant changes in their **(N)** orientations (p<0.01, 2-way ANOVA followed by Dunnet’s multiple comparison test). **(O)** Enthesis strength correlated with BMD (R=0.60, p<0.001), cortical thickness (R=0.69, p<0.001), and trabecular plate thickness (R=0.44, p<0.001). Enthesis toughness correlated with tendon cross-sectional area (R=0.44, p<0.01, Pearson correlation). (* p<0.05, ** p<0.01, *** p<0.001, **** p<0.0001, ANOVA followed by the Dunnett’s multiple comparison test unless otherwise reported).

Regardless of treatment, all specimens failed via avulsion under monotonic loading (Fig. 5B). Healthy and overuse-degenerated attachments failed catastrophically, showing little post-yield behavior, while underuse-degenerated attachments failed at lower forces and showed distinct post-yield behavior (Fig. 5C). Pathologic loading led to distinct changes to enthesis failure pattern. Underuse increased fracture area by as much as 1.9-fold compared to that of control (p<0.01, Fig. 5D). While overuse-degenerated entheses failed primarily with one bony avulsed fragment, failures in underuse-degenerated attachments showed multiple fragments of avulsed bone (Fig. 5E). Overuse and underuse led to a shift in the fracture location: overuse resulted in more failures at the MF-B interface while underuse resulted in more failures at the B-T interface (Fig. 5F). Both overuse and underuse reduced toughness, but via different mechanisms. Overuse did not affect tendon enthesis strength (Fig. 5G) but led to an increase in stiffness (p<0.01, Fig. 5H), resulting in ~30% decrease in toughness (p<0.05, Fig. 5I). In contrast, underuse led to a decrease in strength (p<0.01, Fig. 5G) and a decrease in stiffness (p<0.05, Fig 5H), resulting in a decrease in toughness (p=0.08, Fig. 5I). Hence, Loss in toughness in overuse entheses was associated with reduced displacement at failure, without a change in strength; loss in toughness in underuse entheses was associated with reduced strength at failure, without a change in failure displacement.

To investigate the architectural adaptations underlying these effects, we characterized changes in the bone underlying the tendon enthesis. Bone morphometric analysis revealed that overuse led to up to 9% gain of bone volume in the humeral head (BV/TV, p<0.01, Fig. 5K), while underuse led to up to 24% loss in bone volume in the humeral head (BV/TV, p<0.0001, Fig. 5K) and up to 22% loss of bone mineral density underlying the attachment (BMD, p<0.0001, Fig. 5L). Study of individual trabecula, via three-dimensional segmentation of the trabecular network into rods and plate microarchitectures (*46*), showed that overuse increased the volume of load-bearing trabecular plates (pBV/TV) as much as 22% (p<0.0001), while underuse decreases this as much as 30% (p<0.0001) (Fig. 5M). Overuse increased the thickness (p<0.01) of individual trabeculae 30%, while underuse decreased the number of trabecular plates by 15% (p<0.0001, Fig. S10). The trabecular network of healthy, cage-active control samples had the highest density of trabecular plates oriented at 90°-60° relative to the dominant fiber direction in the supraspinatus tendon, and the lowest density of trabecular rods oriented in this range (Fig. 5N, Fig. S11). With overloading, trabecular plate density increased in 60°-30° (p<0.05) and 30°-0° (p<0.01) ranges, and with underuse, trabecular plate and rod loss occurred uniformly across all directions (p<0.05). These results demonstrate that overuse loading prompted active reinforcement whereas underloading prompted active removal of the trabecular architecture underneath the enthesis. Thus, the architecture of the bony structure at the tendon enthesis oriented to support and share the load into orientations of relatively low enthesis strength and toughness.

To understand which architectural features drove enthesis mechanical behavior, we correlated enthesis failure properties to bone and tendon microarchitecture using Pearson correlation (Fig. 5O and Fig. S12). Enthesis strength correlated strongly with BMD (R=0.60, p<0.001), cortical thickness (R=0.69, p<0.001), and trabecular plate thickness (R=0.59, p<0.001), but not with tendon cross-sectional area. Enthesis toughness correlated strongly with tendon cross-sectional area (R=0.43, p<0.05), and trended with mineralized fibrocartilage volume (R=0.30, p=0.11). These results are consistent with clinical findings that the loss of mineralized tissue at the attachment site correlates with higher rates of re-tearing following surgical repair (*47*).

## Discussion

This study revealed architectural toughening mechanisms at the enthesis, providing guidance for attachment of dissimilar materials (Fig. 6). First, energy storage in a compliant region of the fibrous attachment was protective, precluding fracture of the intricately architectured transitional tissue and instead leading to fracture of more easily regenerated bone. While counterintuitive, a tough, architectured compliant material attaching two dissimilar materials occurs across nature, e.g., in nacre (*48*), tooth enamel (*49*), and some mollusks (*50*). Compliant attachment layers in engineering have also been used in bottom-up and top-down fabrication of architecture materials (*6*), such as PMMAs inserted in between alumina layers (*51*), to absorb energy and channel crack propagation, and polymeric foams inserted into metallic foams (*52*).

**Fig. 6.**
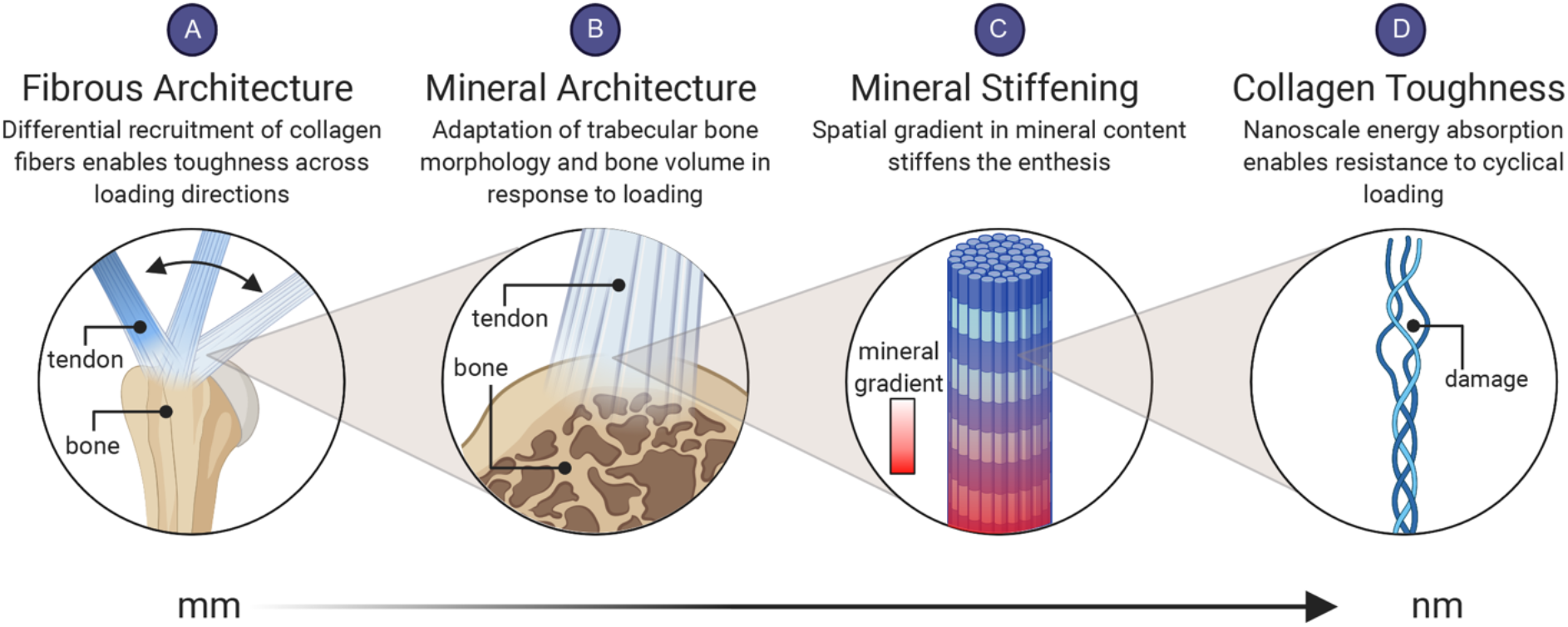
The fibrous and mineral architectures of the tendon enthesis provide multiscale toughening mechanisms for a resilient attachment between tendon and bone. Enthesis toughness is achieved over multiple length scales through unique fibrous and mineral architectures. At the millimeter length scale **(A)** the fibrous architecture of the tendon enthesis allows for fiber recruitment and re-orientation to optimize toughness over strength across a range of loading directions. At the micrometer length scale **(B)** the enthesis actively adapts its mineral architecture to maintain its strength along the axis of loading. At the micrometer-to-nanometer length scale **(C)** a spatial gradient in mineral across the enthesis reduced stress concentrations (*16*). At the nanometer length scale **(D)** collagen damage localization protects against damage prorogating to higher length scales.

Second, the tendon enthesis harnesses its fibrous nature for effective load transfer. Nanoscale energy absorption by collagen molecules resists fatigue loading, while milliscale network behavior enables fiber reorientation, recruitment, and load sharing for toughness across loading directions. A similar concept has been applied to topologically interlocked material panels, with failure shared across contiguous panels and localized to repairable regions (*53, 54*). Distributions of fibers are further optimized at the enthesis to harness the toughness of the entire fibrous network at all loading directions, and to provide enhanced stiffness in the loading conditions for which muscle forces are highest. This relatively simple mechanism provides a principle that can be readily harnessed for engineering.

Additional features of the enthesis that will be more difficult to harness in engineering are compositional adaptions of architecture to physiologic loading. *In vivo* loading models revealed bony architecture actively remodeling to maintain strength along the axis of loading, while compromising overall toughness. Microstructural heterogeneity that toughens fibrous interfaces (*8, 37*) derives in part from mineral nanocrystal reorganization and reorientation (*16*) but controlling these factors, as well as potential mineral binding proteins such as proteoglycans (*55*) and osteopontin (*56*), is currently beyond the scope of current top-down and bottom-up manufacturing techniques. Our findings demonstrated how the tendon enthesis achieves a remarkable balance between strength and toughness through its architecture to resist injurious loads. The toughening mechanisms identified here for the tendon enthesis provide guidance for improving enthesis surgical repair and enthesis tissue engineered scaffolds, as well as approaches for attachment of architectured engineering material systems.

## Materials and Methods

### Sample preparation and study workflow

All animal procedures were approved by the Columbia University Institutional Animal Care and Use Committee. Supraspinatus tendon-to-bone attachment units (humerus-supraspintatus tendon-supraspinatus muscle) were harvested from adult (>12 weeks) male C57BL6/J mice (n = 275). After dissection, samples were fresh-frozen in PBS and stored at –20°C. The experimental workflow was dependent on two categories: (1) unloaded/intact sample characterization (2) loaded sample characterization. For unloaded-sample characterization, defrosted samples were subjected to initial experimental protocol described in the sections below (i.e., secured at appropriate angle of abduction or chemically digested) and imaged via contrast enhanced microCT or via light microscopy, as the imaging techniques were terminal. For characterizing samples undergoing loading, defrosted samples were first scanned by conventional microCT before subjected to experimental protocol and mechanical testing. After mechanical testing, samples were secured at terminal displacements and either submerged in a 5% mercury chloride (HgCl2, Sigma-Aldrich) or fixed with 4% paraformaldehyde (Sigma-Aldrich) to analyze for macroscopic and fiber network level damage or molecular level (collagen) damage.

### Mechanical testing

All samples were mechanically tested in a saline bath at 25°C to prevent thermal collagen denaturation on a table-top tensile tester (Electroforce 3230, TA Instruments) fitted with 10 lb. load cell (TA instruments). Before testing, the supraspinatus muscle was carefully removed from supraspinatus-humerus unit. Samples were placed into custom 3D-printed fixtures (*57*) and supraspinatus tendon were secured between two layers of thin paper (Kimwipe) with a drop of cyanoacrylate adhesive (Loctite, Ultra Gel Control) before mounting onto custom grips. Samples were secured in fixtures and tested in an orientation corresponding to 90° shoulder abduction unless otherwise noted. Specifically, to identify positional contributions to enthesis toughness, samples were fitted to 3D-printed fixtures that secured samples in an orientation corresponding to various angles of abductions (0°, 30°, 60°,120°, n=10 per angle). For all mechanical testing protocols, samples were first pre-loaded to 0.05 N, pre-conditioned by applying 5 cycles of sinusoidal wave consisting of 5% strain and 0.2%/s, and rested for 300 seconds. The unloaded control group consisted of samples that were prepared and mounted in the mechanical tester, but not loaded (n=5).

Quasi-static and monotonic uniaxial loading: post pre-loading, pre-conditioning, and rest, samples were strained in tension at 0.2 %/s to failure (for all loading conditions unless specified otherwise below). The healthy failed control samples (CTRL) were healthy adult enthesis samples strained in tension at 0.2 %/s to failure in an orientation corresponding to 90° abduction. For the interrupted testing, samples were strained in tension at 0.2 %/s to 1 N, 2 N, 3 N (n=3 per rate). To examine the role of strain rate in enthesis failure, samples were tested under three additional strain rates (2 %/sec, 20 %/sec, 200 %/sec, n=10 per rate) until failure. Fatigue loading: after pre-loading and preconditioning, samples were either subjected to 2 Hz sinusoidal loading from 0.1-1 N (1%-20% of failure force, n=4) or 1-3 N (20-70% of failure force, n=5). To investigate molecular level damage localization in the entheses, additional samples were loaded to 10,000 cycles (n=3), 40,000 cycles (n=3), and to failure (> 50,000 cycles, n=5) using the second protocol (20-70% max failure force).

Enthesis structural properties, such as failure load (referred to as strength in text), stiffness, and work to failure (area under the curve through failure load, referred to as toughness in text) were determined from load-deformation curves. Stiffness was calculated by a MATLAB (Matlab2019a, MathWorks) custom algorithm that identifies the best fitting line within a sufficient bin width (i.e., remove data below 10% of max load and above 95% of max load) by implementing the random sample correlation (RANSAC) technique (*58*).

### Contrast enhanced and conventional micro computed tomography (microCT) imaging

Simultaneous visualization of soft and hard tissues of tendon enthesis samples were achieved by staining samples with 5% mercury chloride solution prior to scanning with microCT. A 5% mercury chloride solution was prepared fresh for each experiment day by dissolving Mercury (II) chloride (HgCl2, Sigma-Aldrich) in distilled and de-ionized water (MilliQ water, MilliporeSigma) at room temperature until the saturation was achieved. Tendon enthesis samples, either intact or post-mechanical testing, were submerged in this solution for 24 hours and washed three times in distilled and de-ionized water for 10 minutes each before they were imaged with microCT (Skyscan 1272, Bruker).

We used the same preparations and scan settings when visualizing enthesis samples with both conventional and contrast enhanced microCT. To prepare for scanning, distal end of supraspinatus-humerus unit were embedding in 2% agarose (Sigma-Aldrich) and mounted in the scanning chamber, so that tendon enthesis specimens were hung loosely and in line with the scanning axis. To visualize enthesis samples at specific angles, we used 3D printed fixtures that fixed the samples in the appropriate position when they were mounted in the scanning chamber. Scans were performed with 60kVp, 166uA, and Al 0.5mm filters with isometric resolution of 2.5 μm. To visualize enthesis insertions and failure surfaces, high resolution images were obtained at 0.75 μm resolution, while for whole joint imaging images were obtained at 5 μm resolution. The acquired microCT data were reconstructed with the software (nRecon, Bruker) provided with the CT scanner using alignment optimization and beam-hardening correction. The reconstructed image data was visualized with built-in program (DataViewer and CTvox, Bruker).

### Scanning Electron Microscopy (SEM)

Failed tendon enthesis samples (n=10) were dried at 37 °C, fixed on SEM aluminum pin mounts using carbon tape and silver paint and carbon-coated (30 nm). Prepared samples were imaged by scanning electron microscope (FEGSEM, Quanta 250F, FEI Company, Hillsboro, OR, USA) in backscattered electron mode using a concentric backscattered detector and acceleration voltages of 5-15 KV at a working at different magnifications from 250x to 20,000x. SEM was carried out using facilities at the University Service Centre for Transmission Electron Microscopy, TU Wien, Austria.

### Tendon cross-sectional area, mineralized fibrocartilage area, footprint area, insertion area, and failure area determination

Conventional and contrast enhanced microCT scans of murine tendon enthesis samples were analyzed to determine minimal tendon cross-sectional area, mineralized fibrocartilage (MFC) area, enthesis footprint area, insertion area, and failure area. The minimum tendon cross-sectional area and mineralized fibrocartilage area for each sample was determined from conventional microCT scans that were performed on samples prior to mechanical testing (or prior to staining with HgCl2) and analyzed via built-in image processing algorithms (CTAn, Bruker). Minimum cross-sectional tendon area was determined by thresholding the transverse slices through the tendon, calculating the area encompassing the tendon, and selecting the smallest area of a tendon that is within 500μm from the tendon insertion site. MFC volume was determined by contouring, thresholding, and integrating all the areas of MFC from sagittal slices of humeral head. Since the absorption coefficients of the MFC was in between that of tendon and bone, and did not change significantly between samples, a single range of threshold values was selected to identify and estimate volume of the MFC.

Apparent footprint area, insertion area, and failure area were estimated using HgCl2 stained contrast enhanced microCT images of enthesis samples, as the imaging technique allows for differential absorbance coefficients between each tissue selected. Since the regions of interest were along the curved volume (i.e., humeral head), we developed a custom semi-automated MATLAB (Matlab2019a, MathWorks) routine that calculates the overlapping polyhedron surface meshes from two arbitrary volumes (e.g., humeral head and tendon enthesis) from the same imaging dataset. The first region represents the surface of the bone: either the surface of the humeral head (for calculating footprint area or insertion area), or the surface of avulsed pieces (for calculating failure area). This region was obtained by thresholding and semi-automatically contouring via shrink-wrapping algorithm built-in to the manufacturers’ imaging processing software (CTan, Bruker). The second region for calculating *footprint area* or *insertion area* represents volume of the tendon enthesis that intersects with the surface of the humeral head along the edge of the tendon attachment. The second region for calculating *the failure area* represents a volume that contains only the fractured surface of the avulsed piece. The edges of the second region for in both cases were determined visually by an experienced researcher by manually contouring appropriately slices for each region of interest. The output volume sets were triangularly meshed to determine the surface area between the overlapping volumes.

### Collagen damage visualization

Unloaded and loaded tendon enthesis samples allocated for analyzing molecular-level collagen damage were stained with F-CHP (3 Helix) and visualized via fluorescence microscopy. Post mechanical testing, samples were first secured and fixed at their appropriate displacements with 4% paraformaldehyde (PFA, Fisher Sci) overnight. Tendon enthesis samples were washed 3 times in PBS for 10 min each at room temperature. After washing, each tendon enthesis sample was placed in a tube containing 450 μl of PBS solution. F-CHP staining protocol was adapted from what have described previously in staining rat tendon fascicles (*31*). CF-CHP stock solution (150 μM) was heated at 80 °C for 10 min to thermally dissociate trimeric CHP to a monomeric state and quenched in ice bath for approximately 20 seconds to prevent artificial thermal damage to samples. 50 μl of monomeric CF-CHP were then added to a tube containing tendon enthesis sample, resulting in a final F-CHP concentration of 15 μM. Samples were incubated for overnight at 4°C and washed in PBS 3 times for 30 min in a room temperature to remove any unbound F-CHP molecules. Stained samples were mounted on a glass slide and imaged and captured using an automated ZEISS Microscope (10x objective, excitation at 488nm channel). Images were captured by CCD camera using the built-in image acquisition and stitching features and analyzed with ZEN lite software (ZEISS).

### Positional recruitment model

We consider *N* linear elastic fibers of thickness *t*, each spaced a distance *s* apart, beginning with a fiber that is immediately to the left of a circular bone ridge of radius *R*. When the grip is turned at an angle *θ* to represent positional change, fibers are stretched in that direction. We incorporated three assumptions in building the positional recruitment model as were suggested by the contrast-enhanced imaging results: (1) the outer (bursal side) fibers longer than the inner (articular side) fibers, making the innermost fiber (n=1) shortest; (2) tendon fibers are buckled at high angles of abduction; (3) to simplify, fibers were assumed to be elastic, brittle, and frictionless. During loading, fibers engage, re-orient, and, depending on loading direction, contact its neighbor fibers (or the humeral head) due to curvature of the humeral head (Fig.3a, FigS5). The contact point is determined for each fiber at 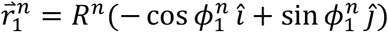, where the radius of the centerline of the wrapped fiber is *R^n^* = *R* + (*n* — 0.5)*t* and the contact angle is 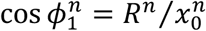. The angle 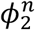 at which contact is lost is determined by the innermost fiber, which always stays in tension. Contact is lost at the point 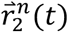 at which the unit vector between 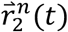 and the connection point on the grip for the strand, 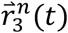, is tangent to the circle formed by the midline of fiber *n*. Using this we can determine the maximum length of a fiber when it is engaged:

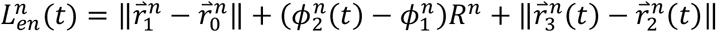

when a fiber is engaged and contact the bone ridge. If a fiber is engaged, but does not contact the bone ridge (when 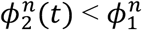):

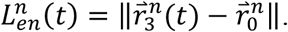

We generated load-displacement curves from this position dependent fiber kinematic model. Expanded details on the positional recruitment model can be found in the supplemental document (Supplementary Text).

### Removal of extracellular matrix components

Glycosaminoglycans (GAGs) from the tendon enthesis samples were chemically digested by adapting a chondroitinase ABC (ChABC) treatment protocol, which is known to degrade GAG chains from tendon (*59*). After conducting a series of concentration and time dependent tests (results not shown), we determined that 0.5 U/mL was an optimum concentration for ChABC for digesting GAGs from tendon enthesis samples. In this protocol, whole samples (humerus-supraspinatus tendon-supraspinatus muscle units) were incubated for 5 days in 2mL of 0.5 U/mL chABC buffered solution (the buffer solution consists of 50 mM Tris, 60mM sodium acetate, 0.02% bovine serum albumin). After 5 days, digested samples were washed in 1xPBS solution 3 times for 30 minutes before subjecting them to microCT imaging and quasi-static mechanical testing. To evaluate the efficiency of ChABC treatment, we performed histological analysis on some samples instead of mechanical testing. These samples (n=2) were fixed in 4% paraformaldehyde for 24 hours, decalcified in formic acid (StatLab, Immunocal), dehydrated in 70% ethanol, and embedded in paraffin. 5 μm thickness paraffin sections were stained with Alcian blue using manufacturers protocol (Alcian Blue Stain Kit, Abcam) and imaged via bright field microscopy with 10× and 40× objectives.

Mineral was chemically removed from the tendon enthesis samples by incubating in 5mL formic acid (Immunocal, StatLab) for 72 hours. Samples were washed in 1xPBS solution 3 times for 30 minutes before subjecting them to microCT imaging to confirm that all the mineral components were chemically digested, and then quasi-static mechanical testing.

### *In vivo* degeneration models

10-week old C57BL6/J mice (n=10/group, Jackson Laboratories) were subjected to two *in vivo* loading models, where the supraspinatus muscle activity was modulated to modify supraspinatus tendon enthesis loading environment. (1) Underuse-degeneration (underuse) was induced via muscle paralysis by bilaterally injecting 0.2 U (0.1U/10μl per 100 g of body weight) of botulinum toxin into the supraspinatus muscles. After injections, mice were allowed to free cage activity for 4 weeks. (2) Overuse-degeneration was achieved using downhill treadmill running (overuse) with an initial rate of 17 cm/s for 10 minutes followed by 25 cm/s for 40 min each day at a decline of 15 degrees, 5 days a week, for 4 weeks (*60*). To acclimate the mice to treadmill exercises, 1 week prior to the overuse protocol, mice underwent training: exercising for each day for 10 minutes at 17 cm/s for 5 days followed by 2 days of rest. For both in vivo models, after 4 weeks since the protocol initiation, mice were euthanized and their supraspinatus tendon enthesis were harvested, soaked in PBS, and stored at −20^0^C.

### Bone morphometry and individual trabecula segmentation (ITS) analysis

Bone morphometry parameters, such as bone volume/total volume (BV/TV), trabecular thickness (Tb.Th.), and trabecular spacing (Tb.Sp.) of the trabecular bone, as well as parameters obtained from ITS analysis were determined using pre-mechanical testing scans of tendon enthesis (5.0 μm resolution). Reconstructed images were first contoured by an experienced user (MG and AA) to only include humeral head proximal to the growth plate as the region of interest (ROI). The ROI were then evaluated using a segmentation algorithm that separates cortical and trabecular bone (CTAn, Bruker). Segmented trabecular images were subjected to subsequent microstructural ITS analysis, where trabecular microstructures were decomposed to individual rod-and-plate-based trabecular microstructural parameters (*46*). In short, the thresholded trabecular bone images were reduced to topology-preserved structural skeletons using digital topological analysis-based skeletonization technique. Each skeletal voxel was then recovered to original topology using an iterative reconstruction method, while classifying whether the resulting trabecular structure belong to either a trabecular plate (surface) or a trabecular rod (curve) using digital topological classification methodology. Microstructural trabecular network and morphology parameters, such as plate-to-rod ratio (PR ratio), rod and plate bone volume fraction (rBV/TV and pBV/TV), number density (rTb.N and pTb.N), and thickness (rTb.Th and pTb.Th) were then evaluated from resultant three-dimensional rod-and-plate classified trabecular morphology. The angular orientational analysis was also performed by evaluating each rod-and-plate angle with respect to perpendicular to the loading axis corresponding to 90 degrees abduction. The average angular distribution for each sample was normalized by the total trabecular volume within each sample’s humeral head.

### Statistical Analysis

Tendon enthesis characteristics, biomechanics results, failure properties, and bone morphometry results were compared between treatment groups using ANOVA and specific differences from control conditions were determined using Dunnett’s multiple comparisons test. P<0.05 was considered significant. Failure properties were correlated to bone morphometry outcomes using Pearson correlation. All statistical analyses were performed using Prism 9 (GraphPad). All data shown as mean ± standard deviation and results from Pearson correlation were expressed using the color map.

## Acknowledgments

The project was funded by National Institute of Health (NIH) U01-EB016422 and R01-AR055580. We acknowledge Fei Fang with the help on *in vivo* degeneration models. We acknowledge Vedran Nedelovski, Amir Davood Elmi and Martin Handelshauser for providing Electron Microscopy images of failed tendon enthesis, which were obtained at the University Service Centre for Transmission Electron Microscopy, TU Wien, Austria.

## Author contributions

M.G., G.M.G., S.T., and V.B., designed the research. M.G., A.C.A., I.K., and B.P.M., developed protocols and performed microcomputed tomography. A.G.S. obtained histological slides. M.G. carried out mechanical testing, confocal microscopy, and contrast-enhanced microcomputed tomography. P.J.T. obtained scanning electron microscopy images. Y.J.H., and X.E.G., performed ITS analysis. M.G., G.M.G., and S.T., analyzed the data and wrote the paper. All authors reviewed and revised the manuscript.

## Data and materials availability

All data is available in the main text or the supplementary materials. The custom codes used in this study, including the positional recruitment numerical model, are available from the authors upon request.

