## Supplemental Materials for "Toughening mechanisms for the attachment of architectured materials: The mechanics of the tendon enthesis"

**This PDF file includes:**

Figs. S1 to S14
Supplementary Text
Table S1
Movies S1 to S4
References (60 to 61)

**Other Supplementary Materials for this manuscript include the following:**
Movies S1 to S4

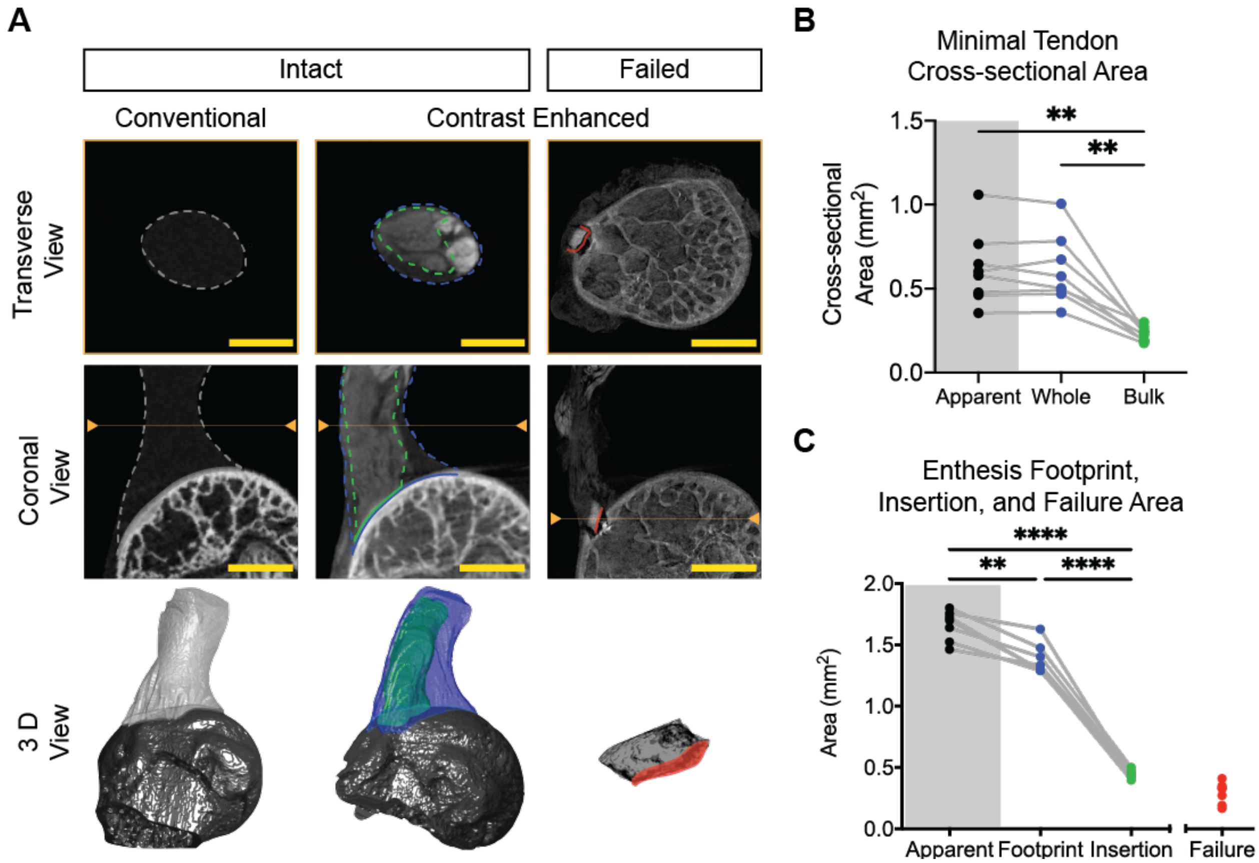


**Fig. S1.**

**(A)** Intact and post-failure images of the tendon enthesis were obtained by conventional and contrast enhanced high resolution micro computed tomography (microCT). The location of the images shown in the transverse view are indicated by the line within the orange arrow heads in the coronal view images. The 3D view was generated using MATLAB from volume contouring and rendering. 2D image scale bars are 800 µm for all images. 3D representations are not to scale. **(B)** Minimal tendon cross-sectional area was obtained from microCT images. "Apparent" area, shown in the gray shaded region, was obtained using conventional microCT; all other measurements were obtained using contrast enhanced microCT. (** $p<0.01$, repeated measures ANOVA followed by the Tukey's multiple comparison test). **(C)** The enthesis footprint, insertion, and failure areas were obtained from microCT images of the endon enthesis. Apparent" area, shown in the gray shaded region, was obtained using conventional microCT; all other measurements were obtained using contrast enhanced microCT. (*** $p<0.001$, repeated measures ANOVA followed by the Tukey's multiple comparison test).

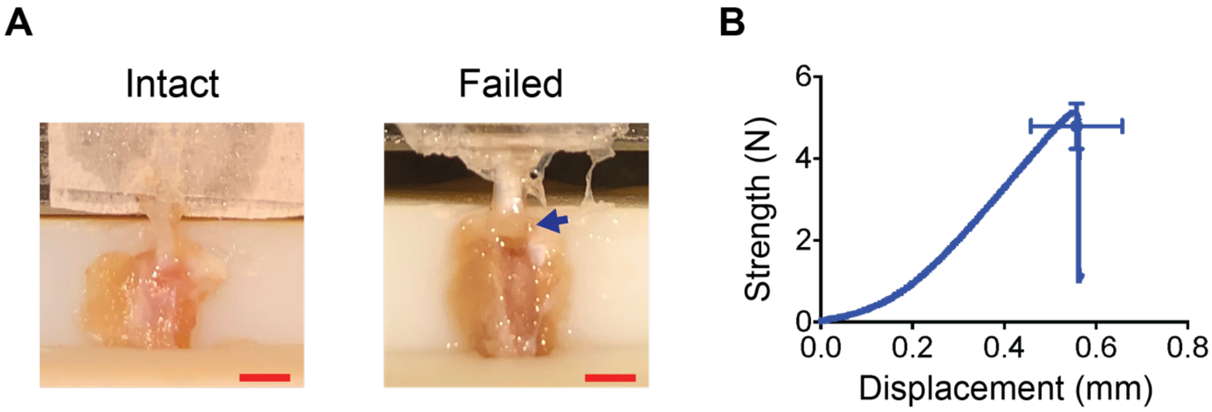


**Fig. S2.**

**(A)** A representative murine tendon enthesis sample is shown before and after mechanical testing (scale bars: 2 mm). While the majority of the primary insertion was avulsed, peritenon tissue surrounding the primary insertion site was still attached post failure (blue arrow). **(B)** A representative tendon enthesis strength (force) vs. displacement curve is shown for a uniaxial quasi-static test to failure. The mean failure strength (force) and failure displacement are represented by the blue dot (cross-head represents standard deviations).

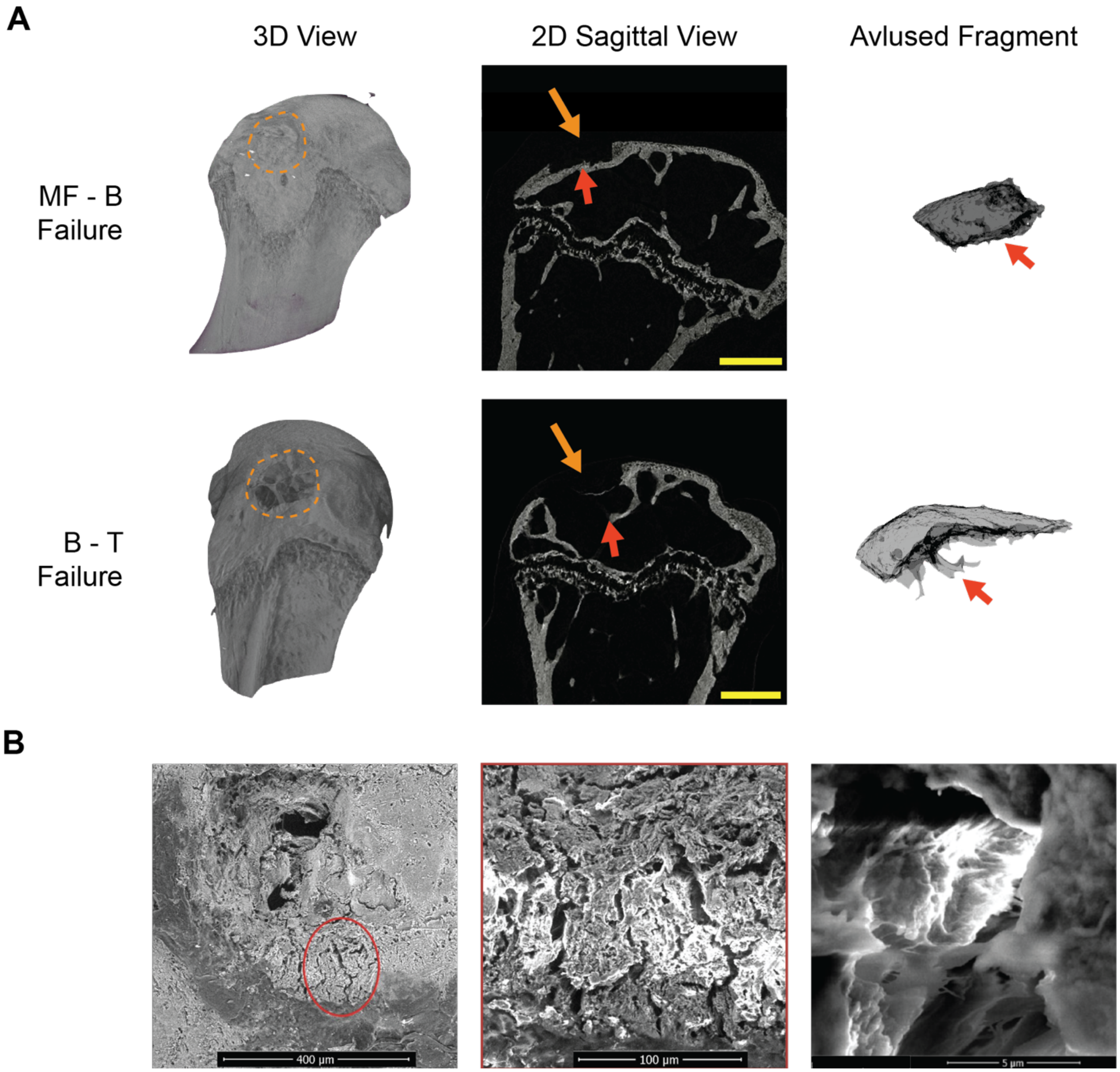


**Fig. S3.**

**(A)** Conventional microCT imaging of post-failure humeral head samples (3D rendering and 2D sagittal view) and 3D visualization of avulsed fragment are shown. Tendon entheses failed either at the interface between mineralized fibrocartilage and bone (MF-B failure type), or in the underlying trabecular bone (B-T failure type) (scale bars: 500 µm). The orange dashed outline and the orange arrows indicate the site of the entheses attachments pre-failure and the humeral head cater post-failure. Red arrows indicate fracture (failure) surfaces. **(B)** Scanning electron microscopy images of failed tendon enthesis samples are shown, focused on the humeral head crater. Crack propagation around the avulsion site was noted. Images were obtained in BSE mode.

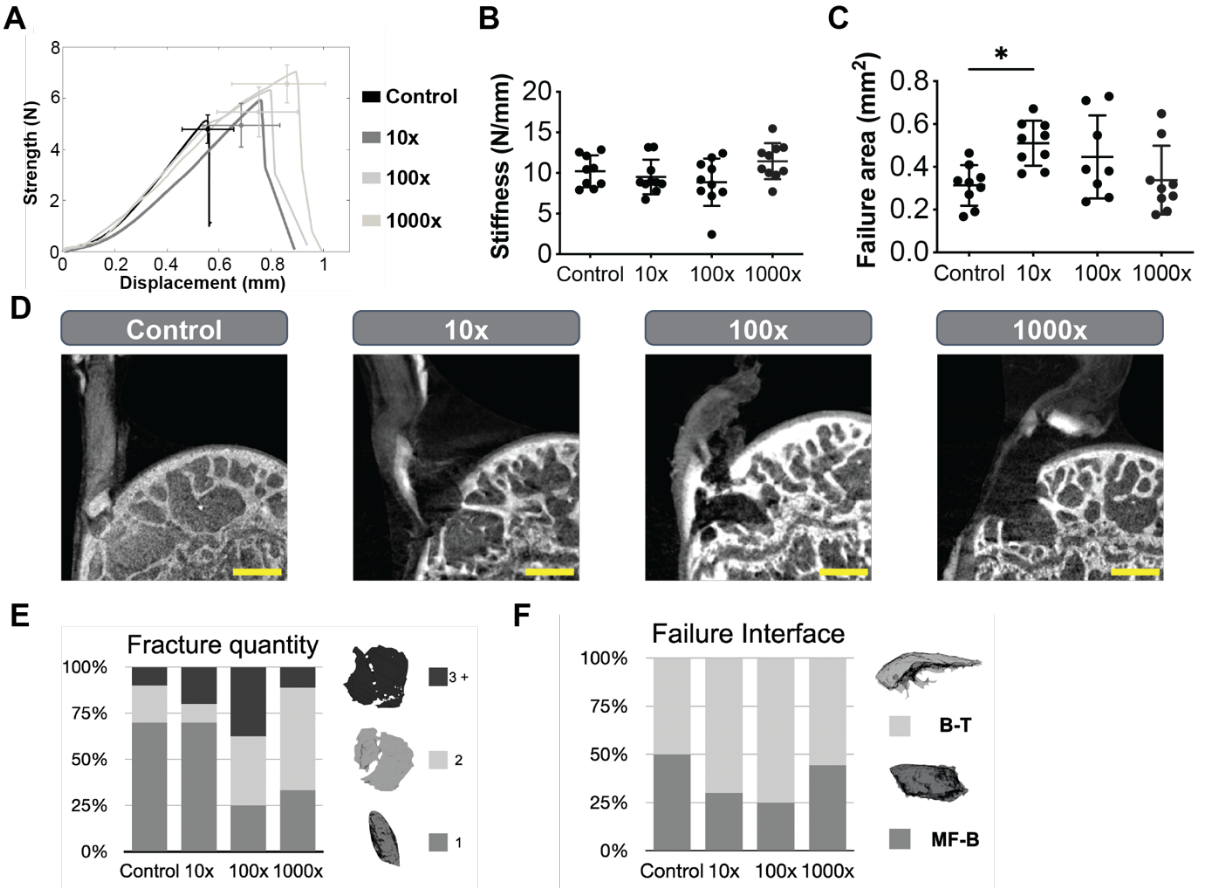


**Fig. S4.**

**(A)** Representative strength (force)-displacement curves for monotonic loadings are shown. **(B)** Stiffness was not affected by increase in strain rates. **(C)**, Avulsed (failed/fractured) area increased with loading rate. (* $p<0.05$, ANOVA followed by the Tukey's multiple comparison test). **(D)** Representative contrast enhanced images of failed samples are shown for all loading rates (scale bars: 250 µm). **(E)** Avulsed (fractured) quantity distribution increased with increasing loading rate. **(F)** Failure interface (MF-B vs. B-T) did not significantly change with increasing loading rate.

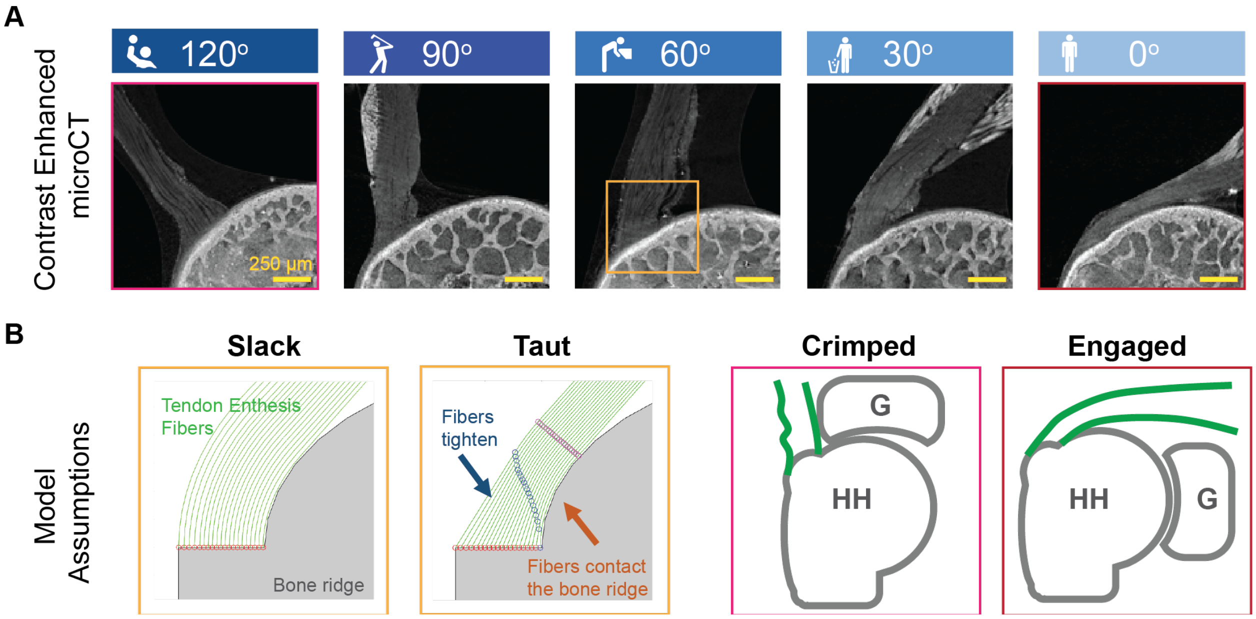


**Fig. S5.**

**(A)** High-resolution contrast-enhanced images of samples are shown at each abduction angle tested (scale bars: 250 µm). **(B)** The positional recruitment model assumed that: (1) fiber length increases with the distance from the bone ridge and (2) fibers get taught once they are engaged (red points to blue points). Depending on loading direction, fibers come into contact with their neighboring fibers or the humeral head and affected by their curvature (blue points to purple points). This allowed for the representation of all fibers as engaged at low angles of abduction (0°-30°) and some fibers intrinsically initially buckled at high angles of abduction (90°-120°). Green: tendon fibers, gray: humeral head bone ridge. G: glenoid, HH: humeral head.

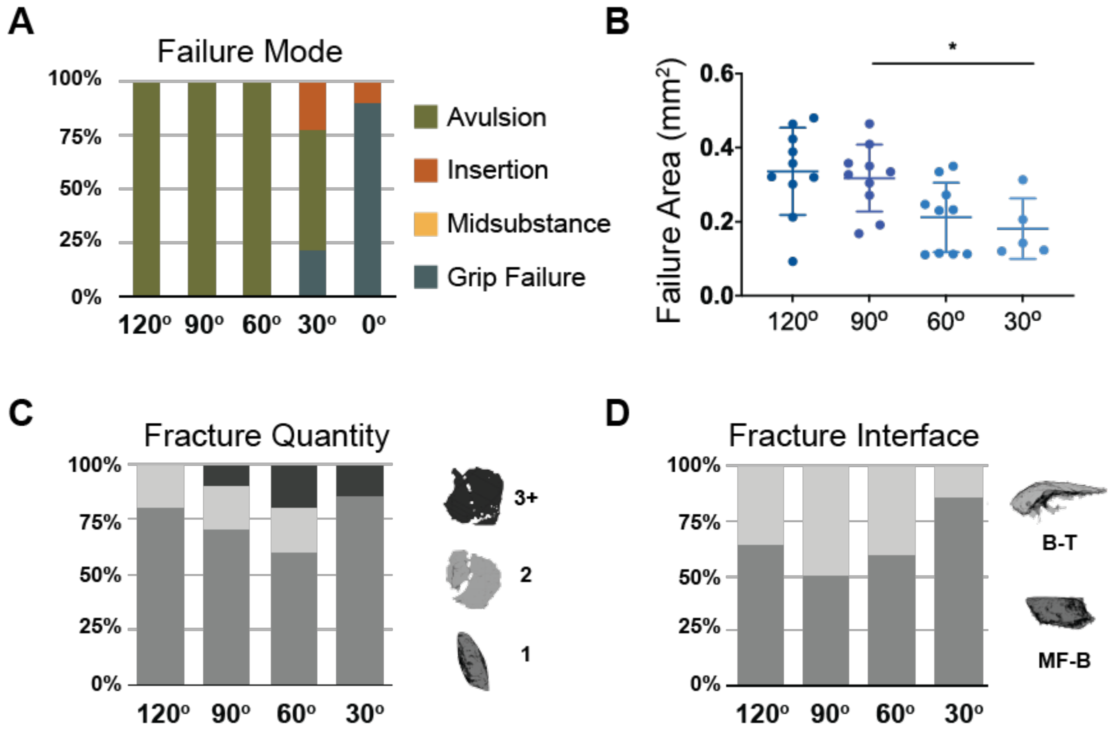


**Fig. S6.**
**(A)** Samples failed primarily via bony avulsion. However, at low angle of abductions (0°-30°), most samples failed at the grips. **(B)** The size of the fractured area decreased at low angles of abduction ($p<0.01$). **(C)** There was a shift towards MF-B type failure when samples were pulled at 30° of abduction. **(D)** Fragment quantity distribution did not show a trend with loading angle.

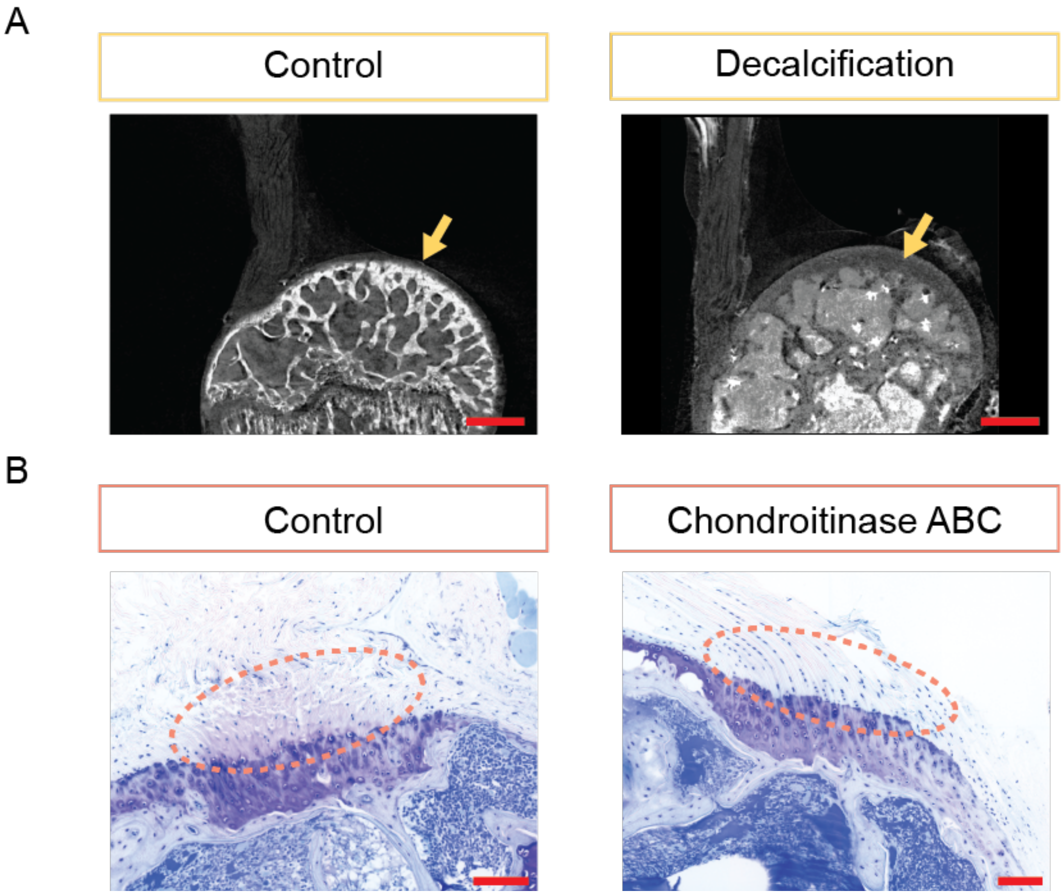


**Fig. S7.**

**(A)** Contrast enhanced microCT imaging showed that formic acid treatment completely removed the mineral from the tendon enthesis and the humeral head bone (scale bars: 500 μm). Yellow arrows indicate changes in the coefficient of attenuation due to the mineral loss in the sample. **(B)** Alcian blue staining of the tendon enthesis showed that chondroitinase ABC treatment removed proteoglycan components from the unmineralized portion of the enthesis (change in staining outlined by orange dashed ellipsoid; scale bar: 100 μm).

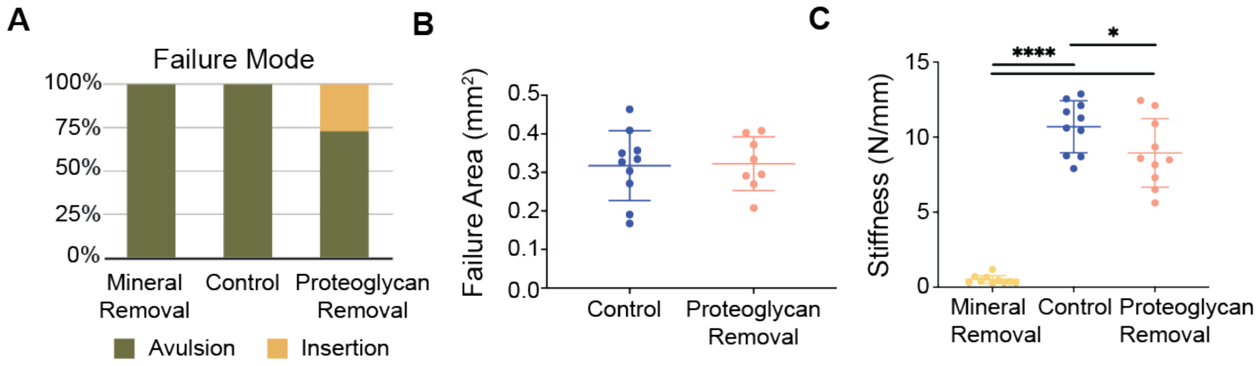


**Fig. S8.**

**(A)** Removal of mineral or proteoglycan did not significantly alter the tendon enthesis failure mode. **(B)** There were no significant differences in failure (avulsed) area due to removal of proteoglycans. Note that the failure (avulsed) area for demineralized enthesis samples was not obtainable with current methodologies. **c,** Removal of mineral or proteoglycan led to significant differences changes in stiffness. (* $p<0.05$, **** $p<0.0001$, ANOVA followed by the Dunnett's multiple comparison test).

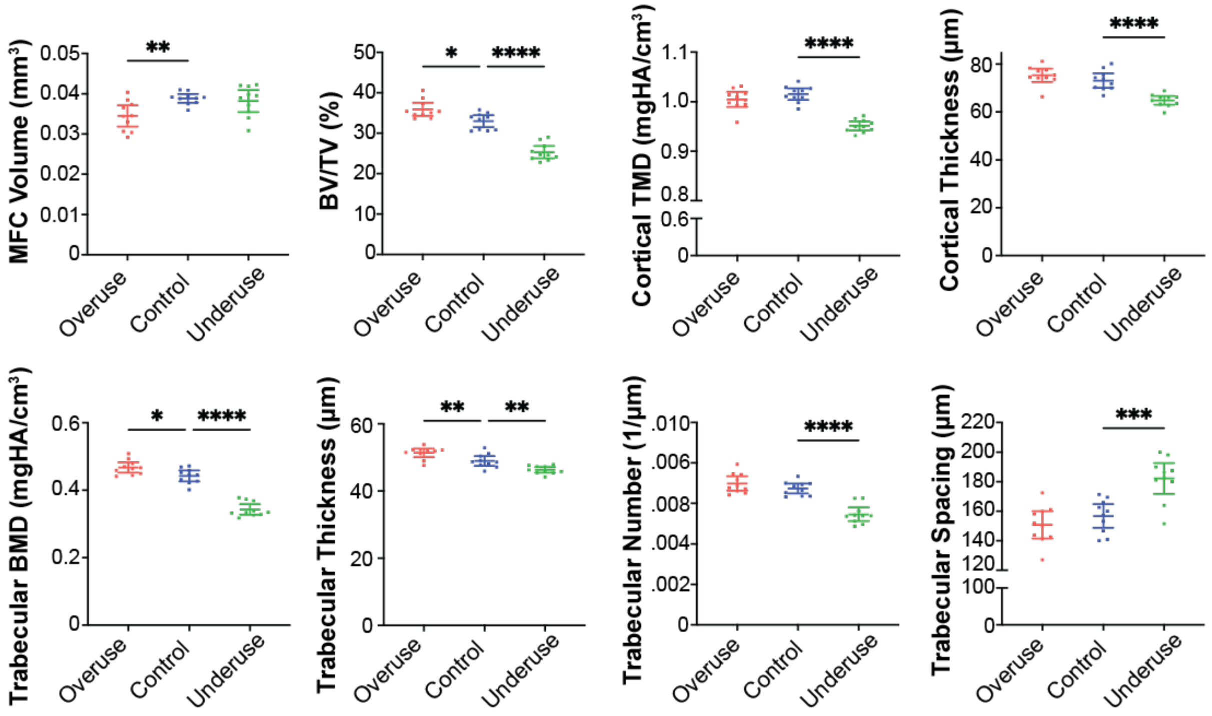


**Fig. S9.**

MicroCT analysis revealed that overuse degeneration led to decrease in mineralized fibrocartilage volume (MFC volume, $p<0.01$). Bone morphometric analysis revealed that underuse led to loss of bone mineral density underlying the attachment (BMD and TMD, $p<0.0001$), reduced cortical and trabecular thickness($p<0.0001$), and trabecular number ($p<0.001$). Overuse degeneration led to gain of bone volume in the humeral head (BV/TV, $p<0.05$), trabecular mineral density (BMD, $p<0.05$), and increase in trabecular thick-ness ($p<0.01$). (* $p<0.05$, ** $p<0.01$, *** $p<0.001$, **** $p<0.0001$, ANOVA followed by the Dunnett's multiple comparison test).

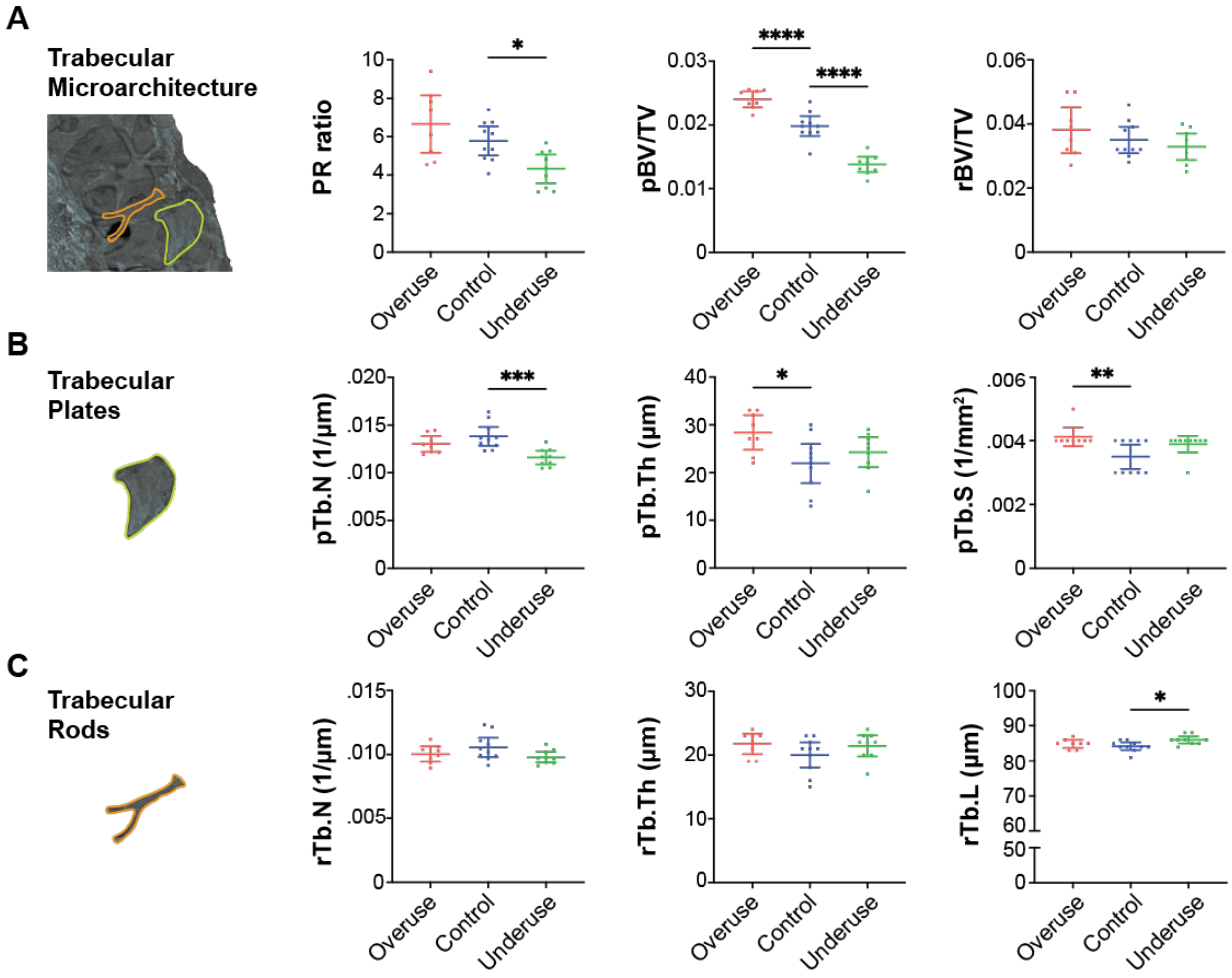


**Fig. S10.**

**(A)** Individual trabecula segmentation (ITS) analysis showed that the ratio between trabecular plates and trabecular rods significantly decreased due to underuse compared to control ($p<0.05$). Trabecular plate volume significantly decreased due to underuse ($p<0.0001$) and significantly increased due to overuse ($p<0.0001$) compared to control. There were no significant differences between groups in trabecular rod volume. **(B)** Trabecular plate analysis showed that the number of trabecular plates (pTb.N) significantly decreased due to underuse ($p<0.0001$), while trabecular plate thickness (pTb.Th) and surface area (pTb.S) significantly increased due to overuse ($p<0.05$). **(C)** Trabecular rod analysis showed that there were no differences in rod number (rTb.N) thickness (rTb.Th). However, the length of trabecular rods (rTb.L) significantly increased due to underuse compared to that of control ($p<0.05$). (* $p<0.05$, ** $p<0.01$, *** $p<0.001$, **** $p<0.0001$, ANOVA followed by the Dunnett's multiple comparison test).

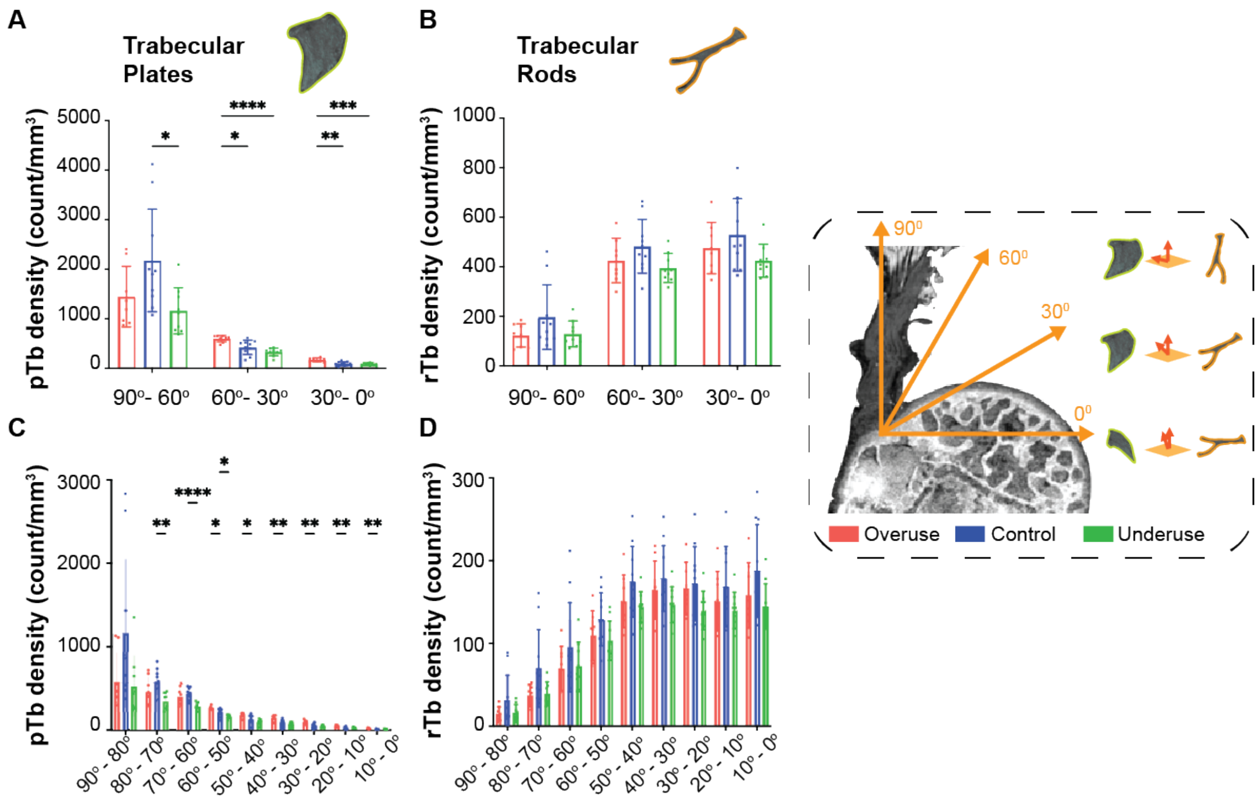


**Fig. S11.**

**(A)-(B)**, Coarse orientation analysis is shown for (A) trabecular plates and (B) trabecular rods**.** **(C)-(D),** Fine orientation analysis is shown for (C) trabecular plates and (D) trabecular rods. The angle was taken with the axis normal to the supraspinatus tendon insertion surface. ( * $p<0.05$, ** $p<0.01$, **** $p<0.0001$, 2-way ANOVA followed by Dunnet's multiple comparison test).

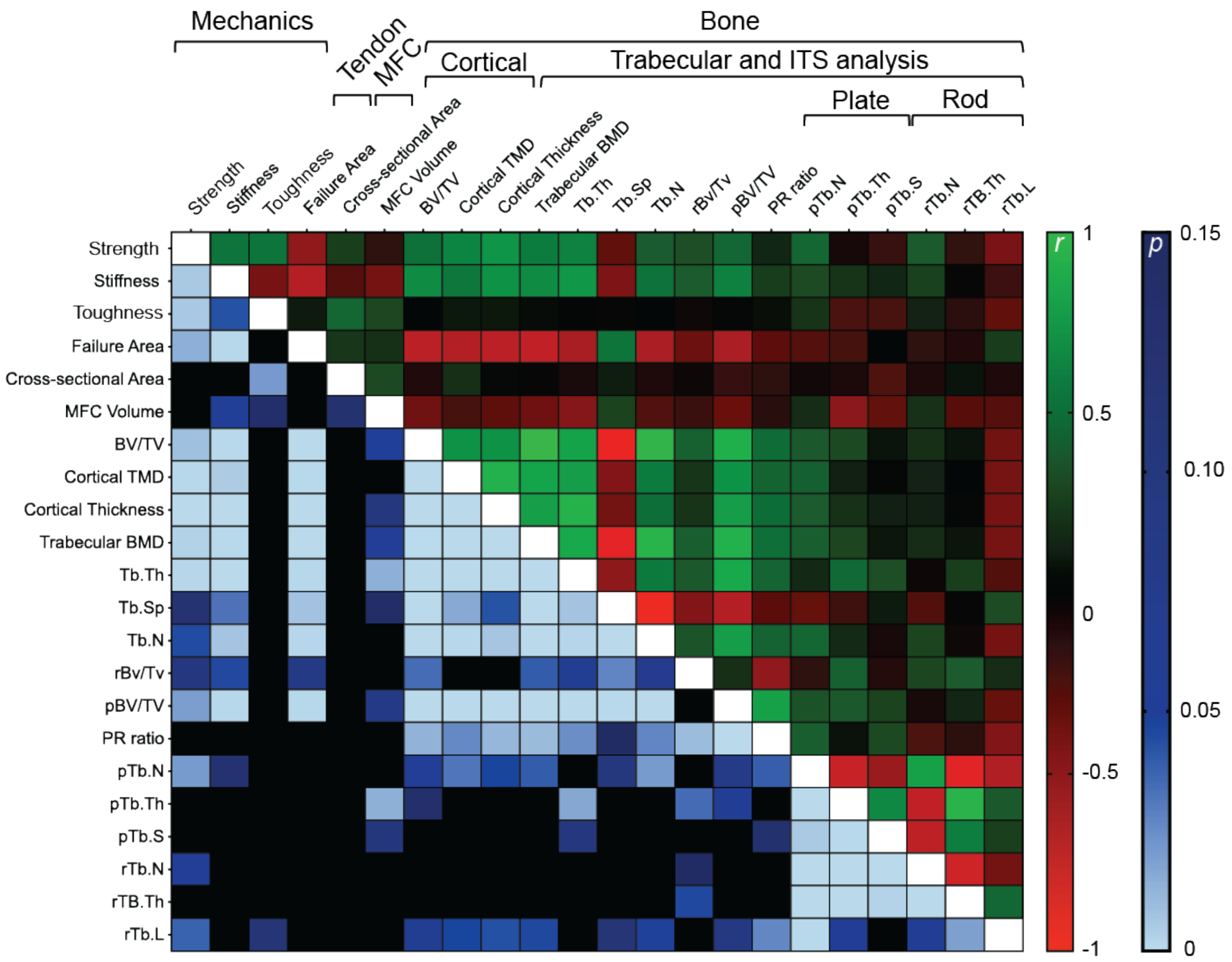


**Fig. S12.**

Pearson correlation results obtained from enthesis mechanical behavior, enthesis failure behavior, enthesis characteristics, and bony architecture under the enthesis. Green represents a positive correlation, while red represents an inverse correlation. Bluegradient show p values

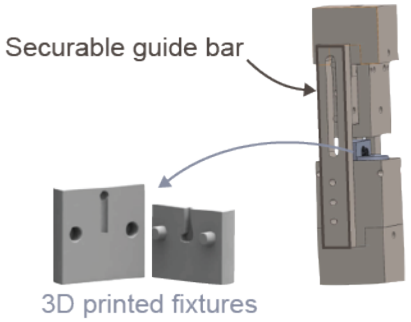


**Fig. S13.**

3D printed fixtures and gripping mechanism used for the mechanical testing protocols.

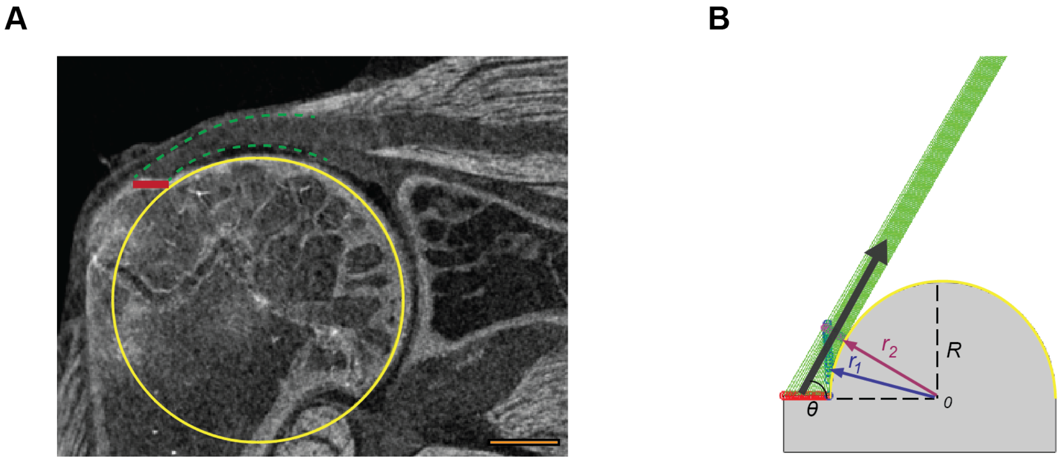


**Fig. S14.**

**(A)** representative contrast enhanced image of the tendon enthesis is shown (scale bar: 500 µm). The orange line indicates the attachment footprint and the yellow circle outlines the circular anatomy of the humeral head. **(B)** A schematic of the positional-recruitment model is shown. Green lines represent enthesis fibers, the gray surface represents the humeral head bone, and the black arrow indicates the direction of the pull. At rest, the outer fibers are longer than the inner fibers. If a fiber is engaged and oriented so that it contacts the bone, it tightens until it reaches the tangent point to the humeral head curvature (blue circles) and the tangent point to the tendon fiber (magenta circles).

**Supplementary Text**

Notes on contrast enhanced microCT scanning and enthesis visualization

While contrast enhancement was non-reversible, the technique was non-destructive; when testing for shrinkage or damage due to processing, the values obtained for whole tendon cross-sectional area for contrast-enhanced samples matched the measurements obtained prior to submerging in mercury (II) chloride solution (apparent cross-sectional area) (Fig. S1). At the site of minimal tendon cross-sectional area, approximately 500 μm above the enthesis, the cross-sectional area of primary tendon fibers was 38% that of the whole tendon (0.23±0.04 $mm^2$), significantly smaller than the primary insertion area (Fig. S1 B, C).

Notes on mechanical testing, enthesis strength, and enthesis toughness

Loading specimens secured with the custom 3D printed fixtures and novel slidable gripping system (Fig. S13) produced force-displacement responses that were highly repeatable (Fig. S2B) and, if needed, interruptible and fixable at prescribed loading/displacement levels. These fixtures also allowed specimens to be recovered post-testing at their failure displacements.

Identifying local strain via conventional optical strain tracking was not possible with the current experimental setup. The bulk tendon is covered by a non load-bearing sheath that does not deform along with the underlying tendon enthesis ( Fig. S1A, blue dotted line). In all failed samples, the sheath was intact post failure (Fig. S2A, transparent tissue). Staining the enthesis samples with Verhoeff's stain to create a speckle pattern in tracking the strain was therefore only able to track sheath surface strain.

In the current paper, we use the nomenclature "strength" (experimentally obtained as a failure load when samples fail) and "toughness" (experimentally obtained as a work to failure, the area under force-displacement curve) to describe the structural properties of mouse entheses. We do not normalize the measurements because: (1) the underlying geometry and local strain of the tendon enthesis could not be defined and (2) tendon entheses did not fail where peak stresses might intuitively be expected (i.e., at the tendon minimum cross-sectional area).

Notes on avulsed (fractured) pieces

To investigate the energy absorption with enthesis failure, we characterized avulsed bony pieces using high resolution microCT. Avulsion area and/or the number of avulsed fragments changed with the loading regime, loading position, and *in vivo* loading conditions. For example, the results for monotonic increases in loading rate showed that the healthy enthesis loaded at high loading rate stored and dissipate enough energy to either propagate its crack to a larger than primary insertion area bony avulsed piece, or from many bony fragments. This failure pattern is consistent with previous observations on rate-sensitivity of bone fracture.

Notes on avulsed (fractured) pieces

The model idealizes the geometry of humeral head (circular bone ridge) and *N* linear elastic fibers with thickness $t$, each spaced a distance $s$ apart make up tendon to bone attachment, beginning with a fiber that is immediately to the left of a circular bone ridge of radius $R$.

The centerline of fiber $n$ inserts into the bone at position:

$$\vec{r}_0^n = -x_0^n \hat{\imath} + 0\hat{\jmath}$$

where $\hat{\imath}$ and $\hat{\jmath}$ are unit vectors parallel to the $x$ and $y$ axes, respectively, shown in Figure 5, and

$$x_0^n = R + \frac{t}{2} + (n-1)s$$

is position of the fiber insertion (red circles at the insertion in the Fig. S5, taut). To note, s>t as s is midline of fiber to midline of next fiber.

We assume that the initial (at rest) length of outer fiber should be larger than inner fiber. The innermost fiber (a fiber that is immediately to the left of a circular bone ridge) is the shortest, and fiber length increases with the distance *s*. This allowed us to represent that: (1) the outer (bursal side) fibers longer than the inner (articular side) fibers, making the innermost fiber (n=1) shortest and (2) tendon fibers are buckled at high angles of abduction. Before defining the initial fiber length $L_0^n$, we derive the posture-dependent fiber engagement theory. If the fiber is engaged and is oriented so that it contacts the bone ridge, then it tightens until it reaches the tangent point (blue circles) to the humeral head curvature (Fig. 2A, Fig. S5, and Fig.S13):

$$\vec{r}_1^{\,n} = R^n(-\cos\phi_1^n\,\hat{\imath} + \sin\phi_1^n\,\hat{\jmath})$$

The radius of the centerline of the wrapped fiber (when the curvature of bony ridge takes an effect) is

$$R^n = R + \left(n - \frac{1}{2}\right)t$$

The contact angle (the angle at which engaged fibers contact the bone ridge and can no longer tighten), is found at

$$\cos\phi_1^n = R^n / x_0^n.$$

The grip holds all fibers in the order they are attached to the bone. When the grip is turned an angle $\theta$ (black arrow) to represent postural change then stretched in a direction $\hat{e}$, the angle at which contact is lost is determined by the innermost fiber, assumed to always stay in tension. Contact is lost at the point $\vec{r}_2^n(t)$ at which the unit vector between $\vec{r}_2^n(t)$ and the connection point on the grip for the strand, $\vec{r}_3^n(t)$, is tangent to the circle formed by the midline of fiber $n$:

$$(\vec{r}_3^n(t) - \vec{r}_2^n(t)) \cdot \vec{r}_2^n(t) = 0$$

or

$$\vec{r}_3^n(t) \cdot (-\cos\phi_2^n(t)\hat{\imath} + \sin\phi_2^n(t)\hat{\jmath}) = R^n$$

Writing $\vec{r}_3^n(t) = r_3^n(e_{3x}^n(t)\hat{\imath} + e_{3y}^n(t)\hat{\jmath})$, $\phi_2^n$ can be solved from:

$$\cos\phi_2^n(t) = \left(\frac{R^n}{r_3^n}\right)\left(-e_{3x}^n + \sqrt{(e_{3x}^n)^2 + \left(e_{3y}^n\right)^2\left(\frac{r_3^n}{R^n}\right)^2 - 1}\right)$$

The grip is placed so that the innermost fiber is not in tension when the tendon is pulled horizontally. Tension starts with $\theta = 0$ and fibers aligned with the $\hat{\imath}$ direction, and with fiber $n$ connected at:

$$\vec{r}_3^n(0^-) = x_3^0\hat{\imath} + R^n\hat{\jmath}$$

where $x_3^0$ is the same for all fibers. The grip is then rotated by an angle $\theta$ about the center of the insertion site, so that:

$$\vec{r}_3^n(0^+) = \mathbf{Q}(\theta)(\vec{r}_3^n(0^-) - \langle\vec{r}_0^n\rangle)$$

where, for evenly spaced fibers, the insertion is centered at the average position $\langle \vec{r}_1^n \rangle = \frac{1}{2}(\vec{r}_1^N + \vec{r}_1^0)$, and the rotation matrix is:

$$\mathbf{Q}(\theta) = \begin{bmatrix} \cos\theta & -\sin\theta \\ \sin\theta & \cos\theta \end{bmatrix}$$

Now, as we previously assumed, the initial (at rest) length of outer fiber should be larger than the inner fiber. The inner most fiber (a fiber that is immediately to the left of a circular bone ridge) is the shortest, and fiber length increases with the distance s, then initial length for n[th] fiber can be expressed as

$$L_0^n(t) = (\phi_2^n)R_0^n + \|\vec{r}_3^n(t) - \vec{r}_2^n(t)\|$$

where $R_0^n = \|\vec{r}_o^n\| = R + \frac{t}{2} + (n-1)s.$

The maximum length of a fiber for it to be engaged when contacting other fibers on the bone ridge is then:

$$L_{en}^n(t) = \|\vec{r}_1^n - \vec{r}_0^n\| + (\phi_2^n(t) - \phi_1^n)R^n + \|\vec{r}_3^n(t) - \vec{r}_2^n(t)\|$$

where $\phi_1^n$ and $\phi_2^n(t)$ are in radians.

Another possibility is that the fiber is engaged but does not contact the fibers around the bone ridge. This occurs when where $\phi_2^n(t) < \phi_1^n$ and:

$$\frac{\vec{r}_3^n(t) - \vec{r}_0^n}{\|\vec{r}_3^n(t) - \vec{r}_0^n\|} \cdot \hat{j} \equiv \hat{e}_{03}(t) \cdot \hat{j} < \frac{\vec{r}_1^n(t) - \vec{r}_0^n}{\|\vec{r}_1^n(t) - \vec{r}_0^n\|} \cdot \hat{j} \equiv \hat{e}_{10}(t) \cdot \hat{j}$$

In this case,

$$L_{en}^{n}(t) = \|\vec{r}_3^{\,n}(t) - \vec{r}_0^{\,n}\|$$

We generated load-displacement curves from this posture depending fiber kinematic model. To simplify fibers were assumed to be elastic, brittle, and frictionless. Strain-displacement relationship were given as

The stretch ration was calculated as follows:

$$\lambda^n(t) = \begin{cases} 1, & L_{en}^{n}(t) \leq L^n(0) \\ \frac{L_{en}^{n}(t)}{L^n(0)}, & L_{en}^{n}(t) > L^n(0) \end{cases}.$$

A linear constitutive law was used for analysis:

$$F^n(t) = \begin{cases} 0 & \lambda^n(t) \leq 1 \\ K(\lambda^n(t) - 1), & \lambda^n(t) > 1 \end{cases}$$

,where K was bulk modulus (stiffness) for the enthesis fibers. The model parameters used for positional recruitment analysis is represented in Table S1.

**Table S1**.

Model parameters used for postural recruitment analysis.

| | parameter | Value |
|---|---|---|
| Number of fibers | $N$ | 20 |
| Fiber thickness | $t$ | 0.01 |
| Distance between fibers | $s$ | 0.015 |
| Humeral head radius | $R$ | 1 |
| Gauge length | $Min\ L^n(0)$ | 2.5 |
| Failure strain | $max\frac{L_{en}^n(t)}{L^n(0)}$ | 0.2 |
| Fiber bulk stiffness | K | 1 |

**Movie S1.**
3D volume rendering of intact mouse enthesis samples. Artificial coloring was added to help visualize the tendon enthesis.

**Movie S2.**
2D image stacks of a representative failed tendon enthesis sample.

**Movie S3.**
3D visualization of a representative failure crater where the tendon enthesis is failed. The images were obtained using high-resolution convectional microCT at sub-micrometer resolution (0.75 μm).

**Movie S4.**
3D visualization of mouse glenohumeral joint visualized using contrast enhanced microCT technique.

**References.**

61. M. M. Panjabi, A. A. White, W. O. Southwick, Mechanical properties of bone as a function of rate of deformation. *J. Bone Joint Surg. Am.* (1973), doi:10.2106/00004623-197355020-00007.

62. T. M. Wright, W. C. Hayes, Tensile testing of bone over a wide range of strain rates:

effects of strain rate, microstructure and density. *Med. Biol. Eng.* (1976), doi:10.1007/BF02477046.
